# A quantitative and predictive model for RNA binding by human Pumilio proteins

**DOI:** 10.1101/403006

**Authors:** Inga Jarmoskaite, Sarah K. Denny, Pavanapuresan P. Vaidyanathan, Winston R. Becker, Johan O.L. Andreasson, Curtis J. Layton, Kalli Kappel, Varun Shivashankar, Raashi Sreenivasan, Rhiju Das, William J. Greenleaf, Daniel Herschlag

## Abstract

High-throughput methodologies have enabled routine generation of RNA target sets and sequence motifs for RNA-binding proteins (RBPs). Nevertheless, quantitative approaches are needed to capture the landscape of RNA/RBP interactions responsible for cellular regulation. We have used the RNA-MaP platform to directly measure equilibrium binding for thousands of designed RNAs and to construct a predictive model for RNA recognition by the human Pumilio proteins PUM1 and PUM2. Despite prior findings of linear sequence motifs, our measurements revealed widespread residue flipping and instances of positional coupling. Application of our thermodynamic model to published in vivo crosslinking data reveals quantitative agreement between predicted affinities and in vivo occupancies. Our analyses suggest a thermodynamically driven, continuous Pumilio binding landscape that is negligibly affected by RNA structure or kinetic factors, such as displacement by ribosomes. This work provides a quantitative foundation for dissecting the cellular behavior of RBPs and cellular features that impact their occupancies.

## Introduction

A grand challenge in biology is to understand, predict, and ultimately control the gene expression programs that allow cells to function. RNA processing is central to regulation of gene expression and includes alternative splicing, 5´- and 3´-processing, nuclear export, cytoplasmic localization, translation, and decay. Each step is regulated by a suite of RNA-binding proteins (RBPs), which constitute >5% of the eukaryotic proteome (Mitchell and Parker, 2014; Muller-McNicoll and Neugebauer, 2013; Singh et al., 2015). By binding specific sequence or structure elements, RBPs can provide coordinated regulation of sets of functionally related RNAs, as shown in early studies for the control of iron metabolism transcripts by iron regulatory proteins, the regulation of functionally distinct mRNA cohorts by individual PUF proteins, and the coordinated binding of synaptic protein transcripts by NOVA (Gerber et al., 2004; Keene and Tenenbaum, 2002; Rouault, 2006; Ule et al., 2003).

Given the central importance of RBPs in posttranscriptional regulation, defining and predicting RBP interactions has been a major research focus (e.g., (Darnell, 2010; Dominguez et al., 2018; Hogan et al., 2008; Hogan et al., 2015; Ray et al., 2013; Riordan et al., 2011; Wheeler et al., 2018)). Transcriptome-wide RNA target sets have been identified for hundreds of RBPs across multiple organisms, facilitating elucidation of RBP roles in numerous regulatory processes (e.g., (Darnell, 2010; Gerber et al., 2004; Guenther et al., 2018; Nussbacher and Yeo, 2018; Ule et al., 2003; Xue et al., 2009)). While current RNA target databases provide immense value, several critical limitations to our current knowledge remain.

First, RBP targets are commonly defined in a binary manner, with RNA molecules considered either ‘targets’ or ‘non-targets’ of a given RBP. However, binding is a continuum, determined by RBP affinities, RBP and target concentrations, and other cellular factors. Therefore, quantitative affinity measurements are needed to define and predict RBP binding occupancies across the RNA sequences present in a cell—i.e., the RBP binding landscape—and the subsequent regulation. Indeed, for transcription factors, thermodynamic models have been shown to be more predictive of binding and effects on gene expression than qualitative, binary models (Foat et al., 2006; Gertz et al., 2009; Le et al., 2018; Riley et al., 2015; Segal et al., 2008; Weirauch et al., 2013; Zhao and Stormo, 2011). A second limitation is that most current approaches are optimized for identifying RBP targets, rather than for quantitative determination of RBP affinities or occupancies. Third, current models of RBP specificity are motif-centric and thus assume energetic additivity (Schneider and Stephens, 1990; Stormo, 2000). The accuracy of such models requires quantitative and comprehensive testing (Zhao and Stormo, 2011).

The above limitations and the importance of regulation by RBPs have sparked a growing interest in developing direct, quantitative genomic-scale approaches for measuring RBP/RNA interactions and affinities. Methods such as MITOMI, HiTS-EQ, HiTS-RAP, RNA Bind-N-Seq and RNA-MaP can provide equilibrium binding constants or apparent affinities (Buenrostro et al., 2014; Jain et al., 2017; Jankowsky and Harris, 2017; Lambert et al., 2014; Martin et al., 2012; Tome et al., 2014). Of these, RNA-MaP and HiTS-RAP, two related techniques that utilize a modified sequencing platform and an array of ∼10^5^ unique immobilized RNA species, eliminate an intermediate capture step that can alter binding occupancies, thereby allowing highly accurate thermodynamic and kinetic binding measurements via fluorescence readout ((Buenrostro et al., 2014; Tome et al., 2014) & *vide infra*). Recent studies have demonstrated the utility of RNA-MaP for systematic investigation of RNA-protein and RNA-RNA interactions and for generation of quantitative energetic models (Buenrostro et al., 2014; Denny et al., 2018; She et al., 2017).

We used the RNA-MaP platform to interrogate the sequence preferences of the human PUF family proteins PUM1 and PUM2 across a diverse designed RNA library. PUF family proteins (Figure 1A) are universal in eukaryotes and have been implicated in regulation of mRNA turnover, transport, translation, and localization (Miller and Olivas, 2011; Quenault et al., 2011). In mammals, PUF proteins play important roles in brain and germline development, regulation of innate immunity and other processes (Bohn et al., 2018; Chen et al., 2012; Gennarino et al., 2018; Gennarino et al., 2015; Kedde et al., 2010; Lee et al., 2016; Liu et al., 2017; Miles et al., 2012; Rodrigues et al., 2016; Tichon et al., 2016; Van Etten et al., 2012; Vessey et al., 2010). Extensive prior biochemical, structural, evolutionary, and in-vivo studies of PUF proteins provide a powerful starting point for our quantitative and systematic dissection of specificity (Figure S1A & references therein) and allow us to pose specific biological, engineering, and biophysical questions.

**Figure 1.**
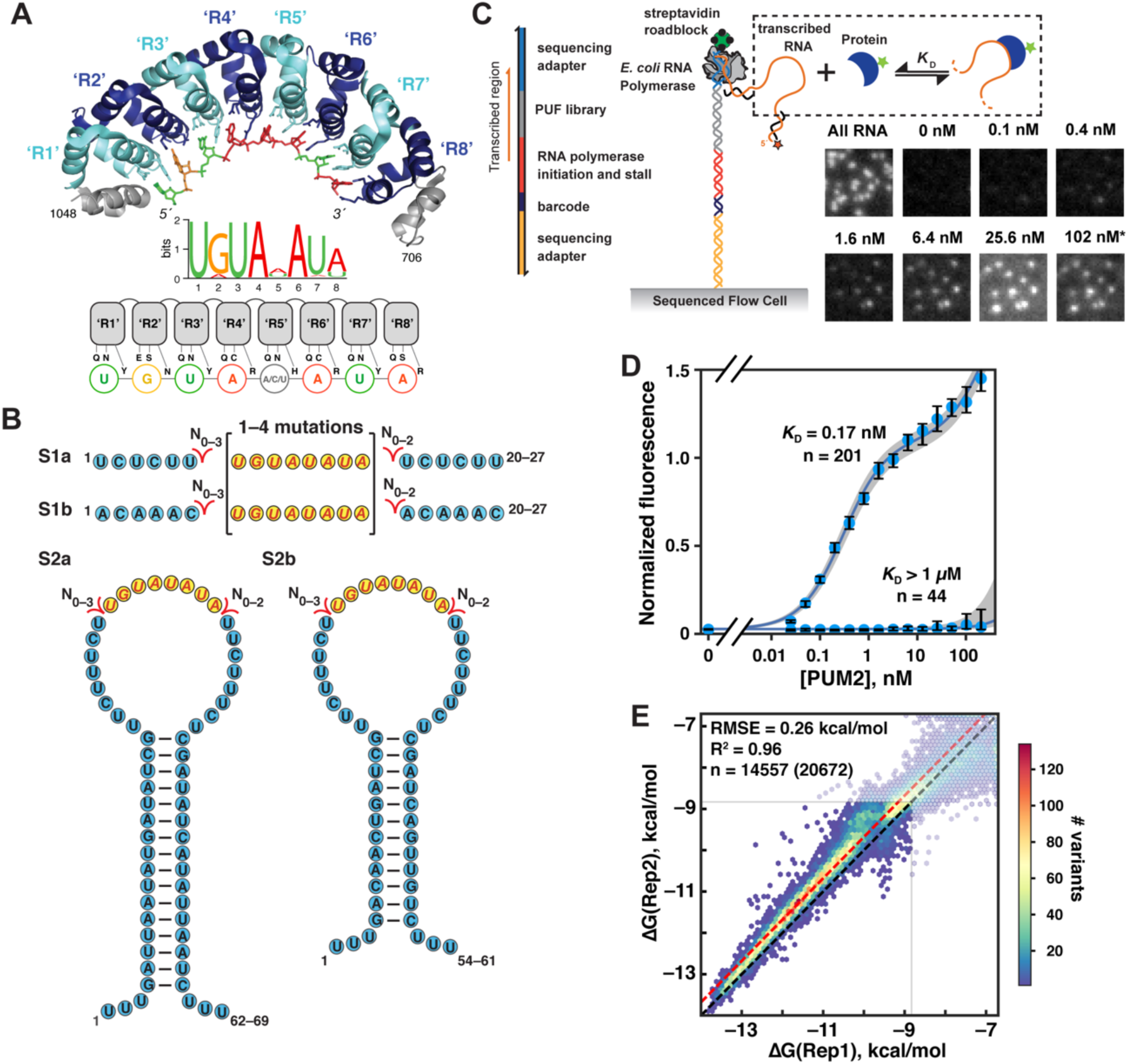
Quantitative High-Throughput Measurements of RNA Binding to PUM2. (A) *Top*: Crystal structure of the RNA-binding domain of human PUM2 bound to UGUAAAUA RNA (PDB: 3Q0Q; (Lu and Hall, 2011)). For simplicity, the eight RNA binding sites (‘R1’–‘R8’) are numbered in the 5′ to 3′ order of bound RNA residues––i.e., the reverse of the order in protein primary sequence. *Center*: Representative PUM2 sequence motif, indicating sequence preferences at each position, as determined by in vivo crosslinking (based on (Hafner et al., 2010); see also Figure S1A). *Bottom*: Schematic representation of PUM2 residues involved in base-specific interactions. The first two residues shown in each repeat form hydrogen bonds and van der Waals interactions with the RNA base, while the third residue stacks with the base (Wang et al., 2002). (B) Scaffolds for studying RNA sequence specificity. Yellow circles indicate the variable region; see Figure S1B for a full description of the sequence variants within this region. (C) *Left*: Schematic representation of an RNA-MaP experiment (Buenrostro et al., 2014) (Methods); see also Figure S1D for DNA library construction and design. *Right*: Representative images of a subset of RNA clusters after incubation with increasing PUM2 concentrations. Asterisk at ‘102 nM’ indicates adjusted contrast relative to other images, due to increased background fluorescence. (D) Representative binding curves for the consensus sequence (UGUAUAUA, S2b scaffold) and a double mutant (<u>C</u>GUAU<u>U</u>UA, S1b scaffold). The number of clusters containing the indicated sequence (n) is noted. Circles indicate the fluorescence in the protein channel normalized by the fluorescence in the RNA channel. Medians and 95% confidence intervals (CIs) across the clusters are shown. Blue lines indicate the fits to the binding model, which includes a non-specific term for PUM2 binding to the PUM2/RNA complex, and the gray area indicates the 95% confidence interval (CI) of the fit (*K*_D_(consensus) = 0.17 nM, CI_95%_ = (0.10; 0.35); *K*_D_(mutant) > 1 μM). (E) Comparison of technical replicates performed on two different flow cells. Data with at least five clusters per experiment and with ΔG error less than 1 kcal/mol (95% CI) are shown. Transparent tiles correspond to ΔG values outside the reliable affinity range (see Methods); ‘n’ corresponds to the number of variants within the high-confidence affinity range, with the total number indicated in parentheses. The black dashed line corresponds to a slope of 1, and the red line is offset by the mean difference between replicates 1 and 2 (0.32 kcal/mol) that accounts for small differences in protein activity and/or dilution. The RMSE was calculated after accounting for this offset (RMSE = 0.42 kcal/mol without accounting for the offset). See also Figure S1.

PUF family proteins have a modular structure of eight conserved tandem repeats that recognize RNA in a sequence-specific manner (Figure 1A) and were chosen for this investigation because this apparent modularity provides a best-case scenario for building a simple predictive thermodynamic binding model (Galgano et al., 2008; Gerber et al., 2004; Miller and Olivas, 2011; Morris et al., 2008; Wang et al., 2002). However, we show that the simplest energetically additive model breaks down and that tight-binding RNA sequences exist that are not represented by previously defined motifs. Our large, quantitative data set enabled the generation of a predictive model for PUM1 and PUM2 binding that includes residue flipping and coupling terms. The model can also be applied to an engineered PUM1 variant, after changing a single parameter to account for the local specificity change. Remarkably, our in vitro-derived binding model *quantitatively* explains prior in vivo crosslinking data (Van Nostrand et al., 2016), demonstrating that RNA binding sites in vivo exhibit, on average, thermodynamically driven occupancies. Our in vitro vs. in vivo analysis further suggests that predicted RNA secondary structures that would inhibit PUM2 binding do not lead to decreased PUM2 occupancy, and thus these structures are likely strongly disfavored in vivo. Our thermodynamic model provides a quantitative foundation for dissecting the cellular behavior of RBPs and represents an early step towards a quantitative and predictive understanding of the complex networks of RBP/RNA interactions and their regulatory consequences.

## Results

### Library design

Starting with the PUM2 consensus motif, which has been determined by pull-down, cross-linking and in vitro selection experiments (Figures 1A, S1A), we designed an oligonucleotide library to systematically address the factors that determine binding specificity (Figure S1B). To control for structural and context effects, each sequence variant was embedded in two to four scaffolds (Figure 1B). We systematically varied the sequence of the PUM2 binding site and the flanking sequence (Figure S1B). We also included insertions to test the potential for noncontiguous binding sites and variants of sequence motifs of related PUF proteins to provide additional sequence variation for testing PUM2 binding models (Figure S1B).

### Massively parallel measurements of PUM2 binding affinities

Using RNA-MaP, we determined PUM2 protein binding affinities for >20,000 distinct RNAs and we report on >5000 herein; sequences designed to address distinct questions will be reported separately. The DNA library was sequenced on an Illumina MiSeq flow cell, followed by in situ transcription in a custom-built imaging and fluidics setup (Figure 1C; (Buenrostro et al., 2014; Denny et al., 2018; She et al., 2017)). The RNA transcripts were immobilized by stalling the RNA polymerase at the end of the DNA template, and RNA-protein association was measured by equilibrating the RNA with increasing concentrations of fluorescently labeled protein and imaging binding to each cluster (comprising ∼1000 copies of an RNA variant) (Buenrostro et al., 2014)(Figure 1C). The resulting binding curves were used to obtain the dissociation constant (*K*_D_) and the corresponding ΔG value (= RTln*K*_D_) of the protein for each RNA variant.

Figure 1D shows representative binding curves for a consensus sequence (UGUAUAUA) and a variant with several mutations (CGUAUUUA) that exhibit divergent affinities (*K*_D_ = 0.17 nM and >1 μM, respectively). For most protein concentrations, protein binding to the consensus sequence followed a canonical binding curve (Figure 1D, UGUAUAUA). At the highest protein concentrations there was a modest additional increase in fluorescence that was only observed for sequences that significantly bound PUM2 and was well fit by a model in which a second PUM2 weakly binds to the RNA/PUM2 complex (Figure 1D, UGUAUAUA vs. CGUAUUUA). This effect was readily accounted for by including a second ‘non-specific’ binding term (Methods) and led to somewhat greater uncertainty in *K*_D_ values for weakly bound RNAs (Figure S1C).

Because our RNA array contained multiple clusters for each sequence variant, numerous binding curves were determined in parallel for each construct. The median number of independent clusters per sequence variant was 23 and 42 in experiment replicate 1 and 2, respectively, greatly exceeding the redundancy of typical biochemical measurements and providing high precision. Molecular variants were included in downstream analysis only when measured in at least five clusters per experiment (Figure S1C), with additional quality filters described in Methods. Independent binding experiments using distinct RNA chips indicated quantitative agreement (R^2^= 0.96; Figure 1E), with average reproducibility within less than two-fold (RMSE = 0.26 kcal/mol) after accounting for a small systematic shift.

### Dissecting and defining PUM2 specificity

PUM1, PUM2 and related Puf3-type PUF proteins appear to recognize RNA in a modular fashion, with each base contacted by one of the eight PUF repeats (Figure 1A; (Wang et al., 2002)). Thus, independent energetic contributions from consecutive RNA bases bound at each of the eight PUF repeats might be expected, as assumed in motif descriptions (Schneider and Stephens, 1990; Stormo, 2000). In this section we test this and other thermodynamic models and answer fundamental questions about PUM1 and PUM2 specificity.

#### Comprehensive analysis of single-mutant variants

We first assessed the binding of all single mutants of the 8mer consensus UGUAUAUA RNA in two to four scaffolds (Figure 2A). At all positions we see the strongest binding for the consensus residue (circled) and we observe very low discrimination at position 5, consistent with prior results ((Dominguez et al., 2018; Galgano et al., 2008; Hafner et al., 2010; Lu and Hall, 2011); see also Figure S1A).

**Figure 2.**
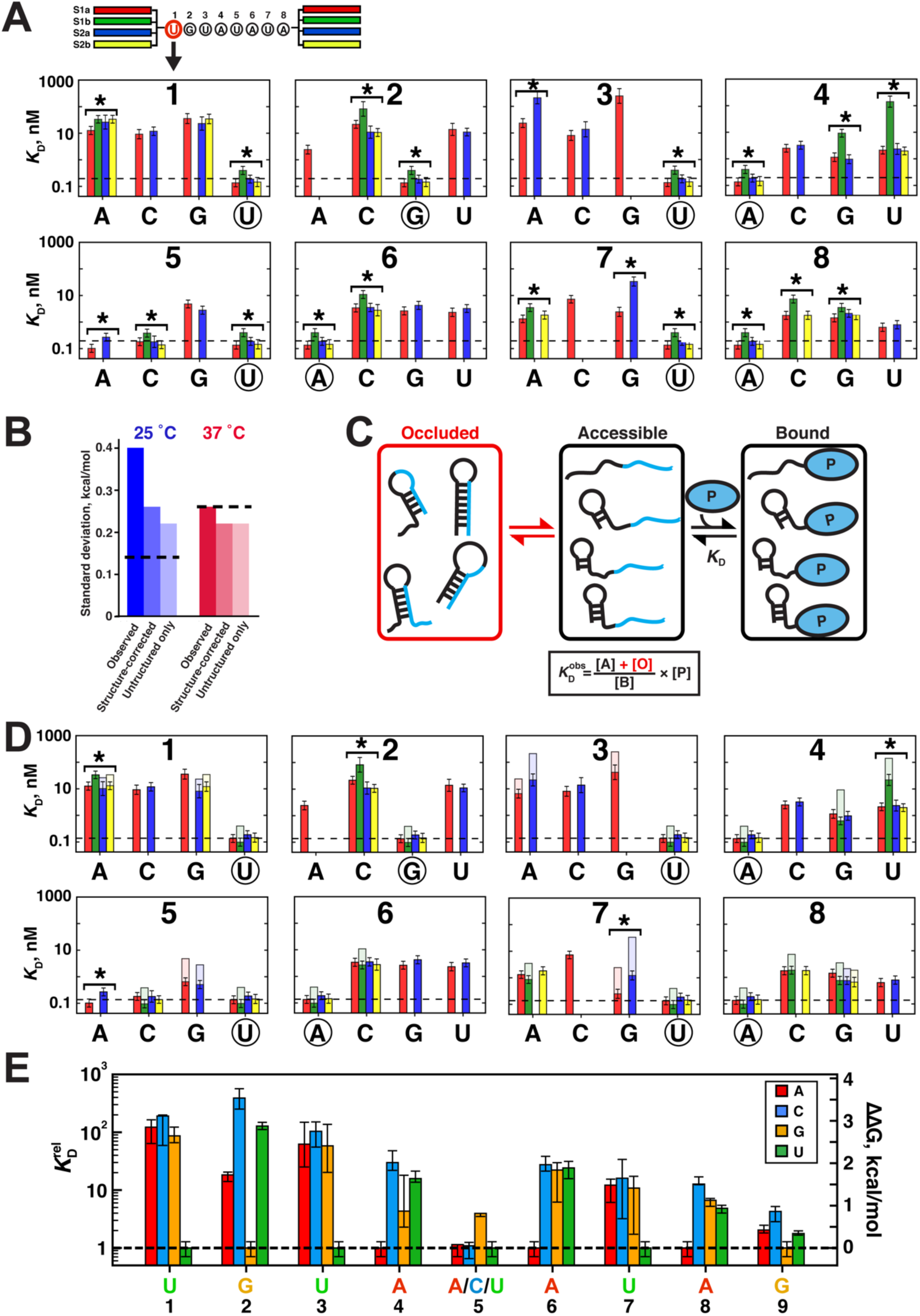
Analysis of Single-Mutant Variant Binding to PUM2. (A) *Top*: Color coding for the scaffolds in Figure 1B; the arrow points to affinities for each position 1 sequence variant. *Bottom*: *K*_D_ values of PUM2 for single mutants at each position of the UGUAUAUA consensus sequence. Bars indicate weighted means of two replicate measurements and error bars indicate weighted replicate errors. The dashed line indicates the average affinity for the consensus sequence across the four scaffolds, and the consensus residue at each position is circled. Asterisks indicate variants with significant differences between scaffolds (10% FDR). (B) Scaffold variance before and after accounting for RNA secondary structure and after excluding sequences with predicted structure. The bars indicate standard deviations of the distribution of differences between each measured value (part *A*) and the scaffold mean for the respective variant; see also Figure S2A and Methods. Dashed lines indicate the standard deviation of measurement error. The experimental standard deviation was higher at 37 than 25 °C because of weaker binding and the absence of an independent duplicate experiment. (C) Model for RNA structure effects on PUM2 binding. Occluded RNA molecules increase the observed dissociation constant (weaken binding) by stabilizing the unbound state (see also Figure S2B & Methods). (D) Single mutant affinities after accounting for structure effects predicted by Vienna RNAfold (solid bars; (Lorenz et al., 2011)); the transparent region indicates the structure correction. Asterisks indicate variants with significant scaffold differences after accounting for structure effects. (E) Summary of single mutation penalties. *Left*: Median effects of each single mutation (residues 1–8) across scaffolds and across 5A/C/U backgrounds at 25 °C, after excluding variants with alternative binding registers and after accounting for structure. Error bars indicate 95% CIs of the median. Mutational effects were calculated relative to the weighted mean affinity for the UGUA[A/C/U]AUA consensus across scaffolds. *Right*: Position 9 specificity, shown relative to the most tightly bound residue (G; see Figure S2D). See also Figures S2, S3.

To assess the robustness of measured single mutant penalties and to identify additional factors influencing PUM2 binding, we compared affinities measured for each RNA variant across scaffolds. While the affinities generally agreed across scaffolds, the spread of deviations was considerably greater than expected from error (Figure 2B; 25 °C, “Observed” vs. dashed line; Figure S2A). Significant deviations between scaffolds occurred in 13 of the 25 sequence variants, at a 10% false discovery rate (FDR; Figure 2A, “*”). Thus, we considered several potential origins of the observed differences between scaffolds.

First, we assessed if RNA secondary structure limits PUM2 access to its site (Figure 2C), to differing extents among scaffolds. If structure affected binding, the differences between scaffolds should decrease at 37 °C. Indeed, smaller differences were observed (Figures 2B, S2A), with only two of the 25 sequence variants exhibiting significant deviations between scaffolds at a 10% FDR (Figure S2C, “*”). Accounting for structure effects with stabilities predicted by Vienna RNAfold (Lorenz et al., 2011) also considerably reduced the between–scaffold deviations (Figure 2B, “Observed” vs. “Structure-corrected”; Figure 2D; the transparent regions denote the structure correction; Methods), with only five of the initial 13 variants with significant inter-scaffold deviations remaining at 25 °C, and none at 37 °C (asterisks in Figures 2A vs. 2D, S2C). A similar decrease in variance was observed after omitting data for RNAs predicted to have significant structure (Figure 2B, “Unstructured only”; ΔG_fold_ > –0.5 kcal/mol). Thus, RNA secondary structure can account for most inter-scaffold variation.

To assess if potential sequence preferences outside the canonical 8mer site affect PUM2 binding, our library included a set of constructs with randomized sequence at the two flanking positions upstream (−2, −1) and downstream (+1, +2) of a common consensus sequence (N = 209 across four scaffolds; Figure S2D). We found modest effects at position +1, with G(+1) bound most tightly (Figure S2D), with no significant effects at other flanking positions, and this G(+1) effect was confirmed in gel shift experiments (Figure S2E). These results identified a new specificity determinant, G at position +1. However, since none of our scaffolds contained a G at this position, flanking effects did not impact the observed differences of single mutant measurements between scaffolds.

Finally, we investigated the possibility of alternative binding registers, which can diminish the observed mutational penalty (Figure S2F). We calculated predicted binding affinities in all possible binding registers along the RNA construct (scaffold + designed binding site) using a model that assumes independent effects of individual mutations (see Methods). Eighteen of our 61 single-mutant measurements across scaffolds in Figure 2A,D had an alternative register with a predicted *K*_D_ within 5-fold of the measured value, which would lower the apparent *K*_D_ by 20% or more (Table S1; Methods). These variants included two of the five mutants that showed significant deviations between scaffolds after structure correction: 1A and 2C (Figure 2D, ‘*’). For the 2C mutant, three of the four scaffolds have alternative registers with affinities matching the observed values (Figure S2G). Thus, in this case it is the seeming outlier (Figure 2D, Scaffold S1b, green) that gives the most accurate mutant penalty. This result underscores the value of multiple scaffolds and the importance of accounting for alternative binding sites for correctly estimating mutational penalties.

To obtain reliable single-mutant penalties despite the alternative register binding, we took advantage of the observations that substituting the uridine at position 5 with A or C residues did not affect the binding affinity (Figure 2D (Lu and Hall, 2011)) and that none of the single mutants in the 5A or 5C backgrounds had stable predicted alternative registers (Figure S3A). We therefore used the median single-mutant effect from the 4–10 constructs per mutation obtained across the 5A, C, or U backgrounds in all scaffolds (Figure S3A), as summarized in Figure 2E. We observed excellent agreement between the values derived from 25 and 37 °C data, with a constant destabilization of binding by (20±10)-fold at the higher temperature (Figure S3A-E). We also observed excellent agreement between gel shift measurements of 14 single mutants and the array measurements (Figure S3F,G).

#### Testing an additive model for PUM2 specificity

If binding of RNA residues by consecutive PUF repeats contributed independently to PUM2 affinity, we should be able to predict PUM2 affinities for our entire library based on additive effects of the measured single mutant penalties (“additive consecutive model”; Figure 3A, *top*). To assess this model, we calculated the predicted affinities for our entire library using 36 terms, one for each residue at each of the 9 recognition sites (8 canonical PUF repeats and the additional G9 site), with the values for these terms determined from our single-mutant data (Figure 2E; Table S2). In the predictions, we accounted for all possible binding registers by calculating the predicted affinity for each 9mer site in a given library variant, followed by calculating the ensemble affinity (Methods). We then compared the predicted and measured affinities for all RNAs in our library with little or no structure (ΔΔG_fold_ > −0.5 kcal/mol; N = 5206). This set included RNAs designed to have two to four mutations throughout the PUM2 consensus sequence, as well as longer sequence variants based on consensus motifs of other PUF proteins and variants featuring insertions and variation in flanking sequence (Figure S1B).

**Figure 3.**
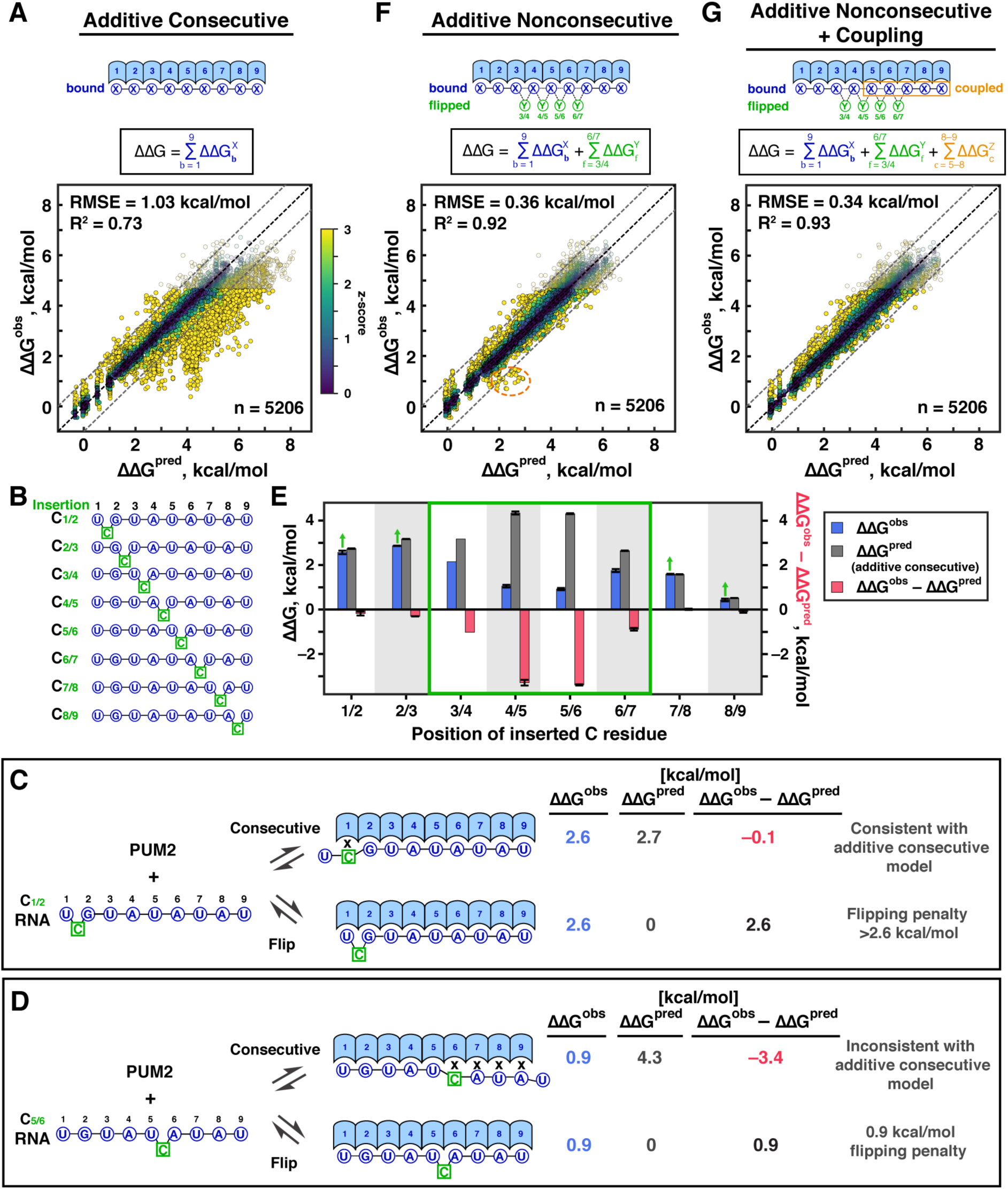
Development of a Predictive Model for PUM2 Specificity. (A) Schematic representation and test of the additive consecutive model. b is the position of bound base and X is the base at position b. 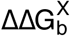 values correspond to the measured single mutation penalties at 25 °C (Figure 2E; Table S2). *Bottom*: predicted versus observed ΔΔG values relative to the UGUAUAUAU consensus sequence for all unstructured variants in our library. Predicted ΔΔG values account for the ensemble of all possible registers along the RNA sequence (Methods). Transparent symbols indicate variants that were bound more weakly than the threshold for high-confidence affinity determination (ΔG > –8.8 kcal/mol); these variants were excluded from determining the R^2^ and RMSE values and from global fitting in parts *F* and *G*. Points are colored based on the deviation from their predicted affinity, divided by the uncertainty of the measurement 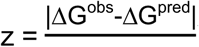. Yellow indicates most significant deviations, and the z-scores were capped at z = 3 for visualization (see also Table S3 for the z-score distribution). The black dashed line is the unity line and the dashed grey lines denote 1 kcal/mol deviation from the predicted value. For direct comparison with parts B and C, data for all unstructured variants (n = 5206) are shown; excluding the single mutants (n = 113) did not significantly alter the fit values (RMSE = 1.04 kcal/mol; R^2^ = 0.73). (B) C-insertion library for base-flipping analysis. (C) Example of an insertion that gives binding consistent with the additive consecutive model. ‘X’ indicates a mismatch. ΔΔG^pred^ corresponds to prediction from additive consecutive model (Figure 3A). With flipping, ΔΔG^pred^ indicates the prediction accounting for bound positions only, which is 0 as the consensus residues are in each site. (D) Example of an insertion that gives binding tighter than predicted by the additive consecutive binding model and provides evidence for base flipping. (E) Summary of observed and predicted ΔΔG values for each of the C insertions in part *A*. Red bars indicate the difference between binding predicted in the best consecutive register for the indicated insertion construct and the observed value. Green box indicates positions at which the observed ΔΔG values are smaller than predicted (negative red bars), suggesting binding with base flipping. Arrows at the other positions indicate that the observed affinities are lower limits for base flipping penalties. (F) Additive nonconsecutive model. Y indicates the residue(s) flipped at position f. Numbering of flipped residues is based on the flanking bound residues; 3/4–6/7 (Figure 3B,E). Model parameters were derived by global fitting and are indicated in Table S4. The dashed orange outline indicates a cluster of outliers with residue coupling (see text). (G) Final model including binding, flipping, and coupling terms. c indicates the positions of coupled residues and Z is the identity of coupled residues. Final model parameters are provided in Table 1. See also Figures S4, S5.

While the predicted and observed binding energies strongly correlated (R^2^ = 0.73), 27% of the observed values deviated from additive predictions by >1.0 kcal/mol, well beyond our experimental error of 0.14 kcal/mol for individual measurements (Figure 3A). The observed deviations were also highly asymmetric, with the vast majority of outliers binding tighter than predicted (Figure S4A). We therefore explored additional features that might lead to the tendency for tighter-than-predicted binding.

#### Residue flipping accounts for most deviations from the additive consecutive model

Several PUF proteins are known to bind RNAs with residues “flipped out” to yield longer, nonconsecutive binding motifs (Gupta et al., 2008; Miller et al., 2008; Valley et al., 2012; Wang et al., 2009; Wilinski et al., 2015), and PUM1, a second human Pumilio protein with >90% sequence identity to PUM2 in its RNA binding domain, has two X-ray structures with bound RNA sequences each with one residue flipped out (Gupta et al., 2008). To assess whether base flipping significantly contributes to RNA binding to PUM2, we had included in our library a set of RNAs with C insertions throughout the UGUAUAUA consensus sequence (Figure 3B), with C insertions chosen because none of the PUM2 repeats preferentially bind C (Figure 2E).

We considered two models for binding of the C-insertion sequences, a null model corresponding to the above additive consecutive model and an extended model that allows for residue flipping. This second model uses the same 36 energy terms for recognition of each residue, but also includes energetic penalty terms associated with flipping out a residue (here the inserted C). Figures 3C and 3D show binding to two RNAs, one that provides evidence for C-flipping (Figure 3D) and one that does not (Figure 3C). In the absence of base flipping, a C insertion would cause one or more mismatches in the PUM2 binding site (“X” in Figure 3C,D), and the relative affinities predicted by the additive consecutive model are indicated (ΔΔG^pred^). For the insertion of C between residues 1 and 2 in Figure 3C, the observed and predicted affinities for a consecutive binding site are the same (ΔΔG^obs^ – ΔΔG^pred^ = –0.1 kcal/mol), providing no indication of base flipping. In contrast, the C insertion between residues 5 and 6 binds 3.4 kcal/mol stronger than predicted by the additive consecutive model, providing evidence for base flipping (Figure 3D). The 0.9 kcal/mol observed destabilization relative to the consensus sequence suggests an energetic penalty for flipping a C residue at this position of 0.9 kcal/mol. Analogous analyses provided evidence for flipping at positions 3/4, 4/5, 5/6, and 6/7 (Figure 3E; green box). Thus, our data suggest that PUM2 can bind RNAs with ‘flipped out’ residues in certain positions, and we modeled this behavior in an extended “additive nonconsecutive model”.

The additive nonconsecutive model combines independent energetic contributions from each of the 9 PUF repeats with the ability to flip up to two residues, each with an associated flipping penalty (Methods). To determine the energetic penalties associated with mutations and with base flipping, this model was fit to our 5206 measured binding affinities (Figure 3F). To avoid overfitting, the parameter values for bound residues (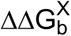 Figure 3) were constrained to the measured single mutant penalties, with an allowed range to account for uncertainty (Figure 2E and Methods). We accounted for all possible binding modes and registers in the global fit (Methods, and see Figure 4). The global fit to the additive nonconsecutive model gave excellent agreement with the data, with RMSE reduced from 1.03 to 0.36 kcal/mol and R^2^ increased from 0.73 to 0.92, relative to the additive consecutive model (Figure 3F vs. 3A). This large improvement is not a consequence of allowing the single-mutant values to vary in global fit, as a global fit to the additive consecutive model with variable single-mutant values gave considerably smaller improvement (Figure S4B,C). Indeed, in the additive nonconsecutive model, 40% of the RNAs are predicted to be predominantly bound with one or more flipped residues. While this model gave nearly symmetric variation around the identity line (Figure 3A vs. 3F; Figure S4A), there remained a subset of outliers (Figure 3F, dashed orange circle) that led us to carry out additional analyses for energetic coupling.

**Figure 4.**
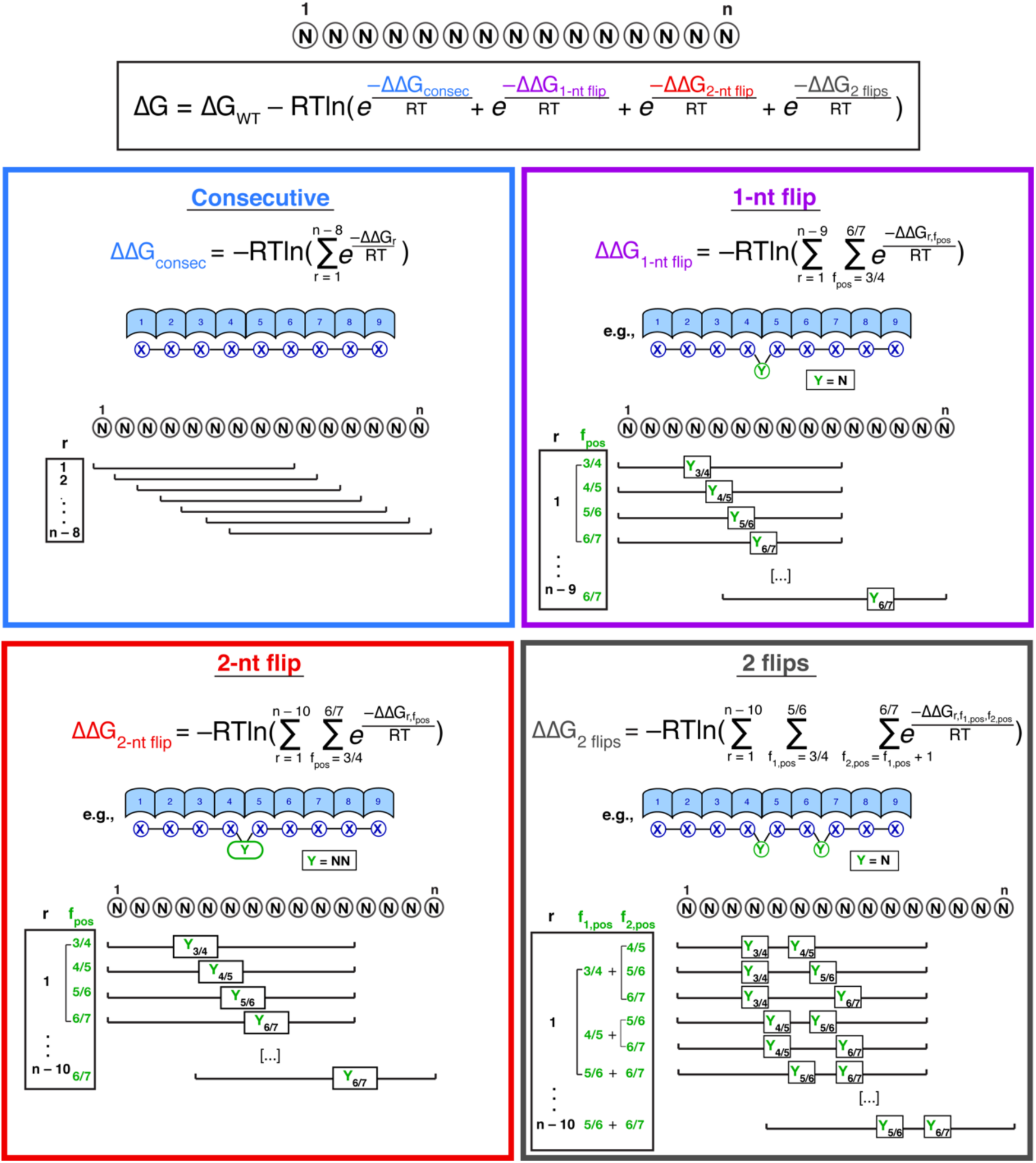
Thermodynamic Model for PUM2 Binding Integrating Binding Modes and Registers. An RNA sequence of length n can be bound in a series of 9–11mer registers (r), within which the RNA residues are variably distributed between bound and flipped positions. Representative subsets of binding registers and base arrangements are shown for each of the four binding modes included in the model: consecutive, 1-nt and 2-nt flips (at a single position), and two flips at different positions. The equations indicate integration of predicted ΔΔG values for all possible binding sites to obtain the final affinity. The ΔΔG values for individual binding site configurations are given in Table 1. ΔG_WT_ is the affinity for the consensus sequence.

#### Energetic coupling between neighboring residues

Inspection of the cluster of variants that bound tighter than predicted even after accounting for flipping (Figure 3F, dashed outline) revealed an enrichment for variants with a G mutation at position 7 accompanied by mutations at position 8, suggesting potential coupling between neighboring mutations. For an unbiased assessment of coupling, we considered all double mutants and plotted deviations of observed effects from the predicted additive effects of individual mutations (Methods; Figure S4D). Coupling between positions 7 and 8 was the strongest, and deviations from additive predictions at all other positions were <0.5 kcal/mol. Analysis of energetic coupling between positions 7 and 8 revealed that: (1) Coupling between positions 7 and 8 occurred with G or C at position 7; (2) For 7G, binding was tighter than predicted (i.e., coupling occurred) only when position 6 was the consensus residue (A) and a pyrimidine was present at position 5 (Figure S4E); these variants fully explained the cluster of outliers observed in Figure 3F (Figure S4E, *top*). (3) For 7C, the base identity at positions 6 and 8 mattered, with deviations from additivity greatest when position 6 was mutated (i.e., not A) (Figure S4F).

We also observed small deviations from additivity at positions 8 and 9 (Figure S4D) and asked whether the small deviations might be real given the overall weak effect from G at position 9 (Figure 2E). Physically, an absence of stable binding in the PUM2 site ‘8’ would be expected to increase the entropic penalty for forming the site ‘9’ interaction and thus might weaken or eliminate this interaction. Indeed, we found that the modest stabilizing effect of 9G relative to other residues was only present with the consensus A at position 8 (Figure S4G). We therefore included this coupling term in our final model.

Figure 3G shows the global fit to our final model that includes additive terms for bound and flipped residues, and the three coupling terms described above (Figure 3G; red boxes in Figure S4E–G). The model accounted for 99% of the data within 1 kcal/mol and gave a slight overall improvement relative to the additive nonconsecutive model (Figure 3F vs. 3G).

#### Evaluating the final PUM2 binding model

To assess the robustness of the final model we performed a series of control fits and analyses (Methods), demonstrating that the fit model parameters were stable to variation in initial parameter values, data resampling, and the use of different fitting methods. Training and testing sets gave essentially identical R^2^ and RMSE values, suggesting that the model was not overfit.

To determine how well the data and model define the individual free energy terms (Table 1), we individually varied each parameter and determined the sensitivity of the RMSE to this variation (Figure S5A–C). As expected, the free energy terms for the consensus residues were highly constrained, as these residues were present in the majority of the 5206 RNAs (Figure S5A, shaded in blue). Varying the penalties for mismatched bound residues gave shallower changes in RMSE, as expected given their lower representation, and, again as expected, this shallowness was most pronounced for the highly destabilizing 5′ mutations that were only weakly represented in the preferred binding registers (Figure S5A).

**Table 1.** **Thermodynamic Parameter Values for Additive Nonconsecutive Coupling Model**

| **A. Term I** | **$\Delta\Delta G_b^X$ (kcal/mol)** |  |  |  |
| --- | --- | --- | --- | --- |
| **Bound residue position b =** | **X =** |  |  |  |
|  | **A** | **C** | **G** | **U** |
| **1** | 3.08 | 2.91 | 3.04 | 0.00 |
| **2** | 1.93 | 3.14 | 0.00 | 3.14 |
| **3** | 2.39 | 2.49 | 2.92 | 0.00 |
| **4** | 0.00 | 1.92 | 1.71 | 1.46 |
| **5** | −0.03 | 0.17 | 0.79 | 0.00 |
| **6** | 0.00 | 1.83 | 1.82 | 1.49 |
| **7** | 1.55 | 1.78 | 1.59 | 0.00 |
| **8** | 0.00 | 1.57 | 1.52 | 1.01 |
| **9** | 0.30 | 0.29 | −0.07 | 0.00 |

| **B. Term II** | **$\Delta\Delta G_f^Y$ (kcal/mol)** |  |  |  |  |  |
| --- | --- | --- | --- | --- | --- | --- |
| **Flipped residue position f =** | **Y =** |  |  |  |  |  |
|  | **A** | **C** | **G** | **U** | **NN\*** | **∅** |
| **3/4** | >2** | 1.79 | >1.5 | 1.41 | >2.5 | 0 |
| **4/5** | >2 | >3 | >2.5 | >2.5 | >2.5 | 0 |
| **5/6** | 1.22 | 1.05 | 1.57 | 0.81 | 2.18 | 0 |
| **6/7** | >2 | 1.77 | >2 | 2.02 | 2.04 | 0 |

| **C. Term III** | **$\Delta\Delta G_c^Z$ (kcal/mol)\*\*\*** |  |  |  |  |  |  |  |  |  |
| --- | --- | --- | --- | --- | --- | --- | --- | --- | --- | --- |
| **Coupled residue positions c =** | **Z =** |  |  |  |  |  |  |  |  |  |
|  | **5** | **6** | **7** | **8** | **6** | **7** | **8** | **8** | **9** | **All other** |
|  | C/U | A | G | C/G/U | C/G/U | C | C/G/U | C/G/U | X |  |
| **5–8** | −1.53 |  |  |  | − |  |  | − |  | 0 |
| **6–8** | − |  |  |  | −0.91 |  |  | − |  | 0 |
| **8–9** | − | | | | − | | | −$\Delta\Delta G_9^{X****}$ | | 0 |

\* Two-nucleotide flip of any sequence.
\*\* “>” Indicates a lower limit (see Figure S5C).
\*\*\* Coupling terms are defined as combinations of residues that meet all of the indicated conditions at the indicated sets of positions. E.g., the coupling term $\Delta\Delta G_{6-8}^Z$ has the value of −0.91 kcal/mol if position 7 residue is a C and position 6 and 8 residues are not A (C/G/U); for all other combinations of sequences, the value of the coupling term is 0 (“−”). See Methods and https://github.com/pufmodel for full description of the model.
\*\*\*\* Coupling term indicates that the position 9 binding term ( $\Delta\Delta G_9^X$ ; Table 1A) is only implemented when position 8 is the consensus residue A.

The fit values for six of the 36 parameters for individual bound residues fell at the limit of the range to which the parameter values were constrained based on the estimated error in single mutant measurements. Removing the constraints for these six parameters gave optimal values within 0.0–0.2 kcal/mol of the previously allowed range of values (Figure S5B). The small deviations could originate from small RNA structural effects or additional weak couplings.

About half of the flipping terms in the model (Figure 3G,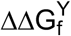 were well constrained, while the other half provided lower limits for the free energy penalties, generally because the penalties were sufficiently high that either no binding or binding in an alternative register was observed (Figure S5C). In addition, the lack of specificity for the ‘bound’ base at position 5 limits the ability to distinguish flipping at position 4/5 versus position 5/6.

#### Implementation of the predictive model of RNA binding by PUM2

To apply our thermodynamic model for PUM2 binding to any given RNA, we considered all possible binding modes to derive a binding ensemble, as illustrated in Figure 4. For the 15 nucleotide RNA example in Figure 4, there are seven possible consecutive (9mer) registers; six 10mer registers, in which a single base is flipped at one of the four possible flipping sites (24 flipped possibilities total); and five 11mer registers that can be bound with two residues flipped at one of four sites (20 possibilities) or with two single residues flipped at two different sites (30 possibilities). For each of the registers, the model calculates individual ΔΔG values by adding the respective individual ΔΔG terms for bound, flipped and coupled residues (Figure 4 and Table 1). The final predicted affinity is then computed from the ensemble of all possible binding site configurations, assuming one bound protein monomer, as specified by the equation in Figure 4. Overall our final binding model is well constrained by the large amount of quantitative data used to build and test it, and the model is complete in that it predicts PUM2 affinity for *any* RNA sequence.

### Evaluating specificity across human Pumilio proteins

Human PUM1 shares 91% sequence identity and 97% sequence similarity in its RNA-binding domain (RBD) with PUM2, and all of the RNA-interacting amino acids are identical between the two proteins (Figure S6A). Prior studies revealed nearly identical RNA sequence motifs and considerable overlap in apparent targets, highlighting the question of why humans retain two seemingly redundant proteins (Figure S1A)(Bohn et al., 2018; Campbell et al., 2012; Dominguez et al., 2018; Galgano et al., 2008; Hafner et al., 2010; Morris et al., 2008). To test potential quantitative differences in PUM1 and PUM2 sequence specificity and to assess if our PUM2-derived binding model could be extended to predict PUM1 binding, we compared PUM1 and PUM2 binding across our RNA sequence library.

PUM1 and PUM2 binding showed remarkably high agreement across the 12,285 sequences for which high-confidence data were obtained (Figure 5A; R^2^ = 0.95, RMSE = 0.24 kcal/mol), indistinguishable from the concordance between PUM2 replicates (Figure 1E). Therefore, our model derived from PUM2 data can also be used to predict PUM1 binding (Figure 5B). Our observation that RNA binding domains of the human Pumilio proteins display identical RNA sequence specificities suggests that any functional differences are fully determined by other factors. Indeed, there is evidence that PUM1 and PUM2 are differentially expressed and diverge in their sequence outside the PUF RBD, allowing for different modification patterns, and potentially different protein interaction partners and differential subcellular localization (Bohn et al., 2018; Kedde et al., 2010; Lee et al., 2016; Liu et al., 2017; Nagaraj et al., 2011; The UniProt, 2017; Van Etten et al., 2012).

**Figure 5.**
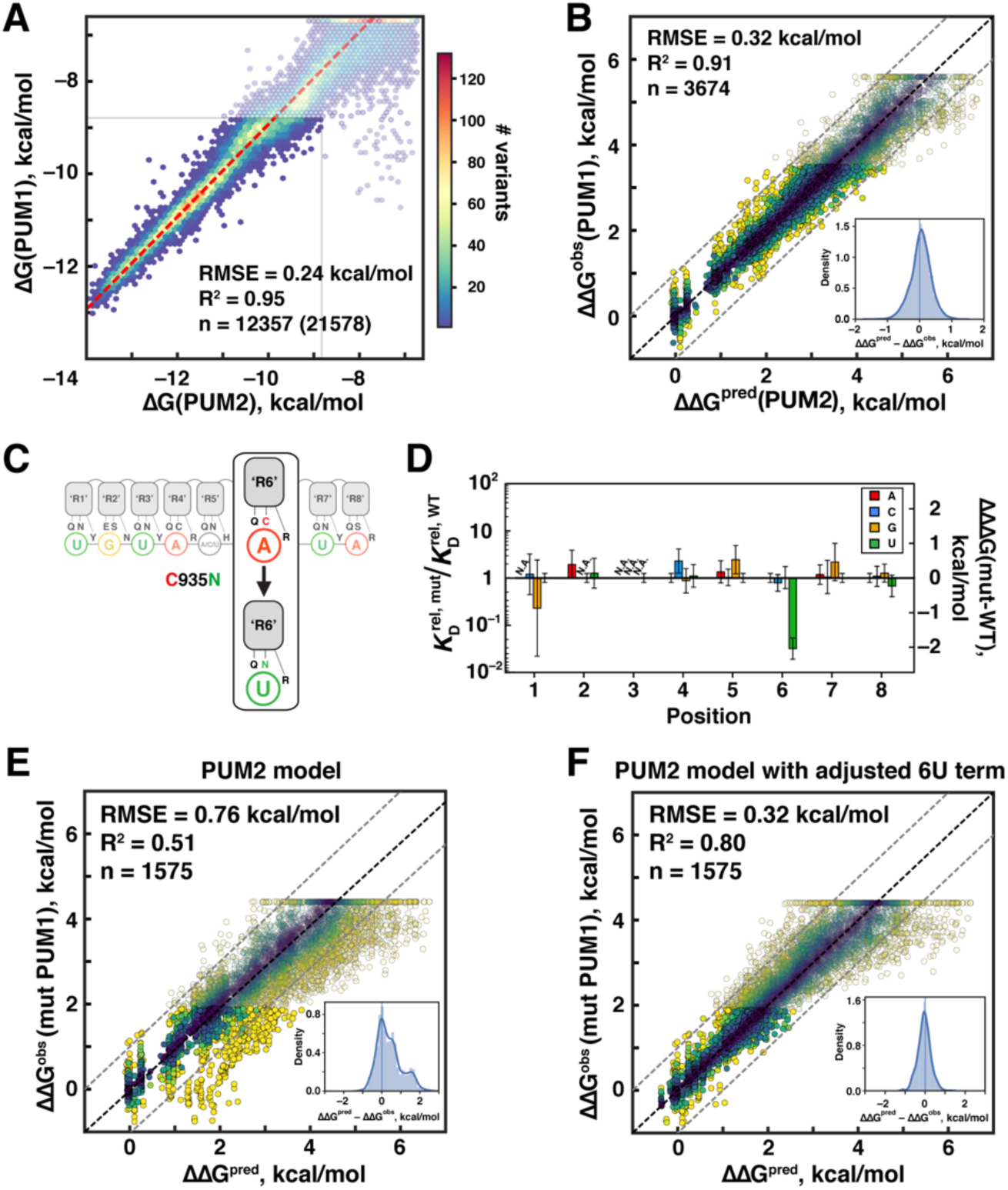
Comparison of RNA Binding Specificities of PUM2 and Wild-Type and Engineered PUM1 Proteins. (A) Correlation between PUM1 and PUM2 affinities across the library. The red line has a slope of 1 with an offset of 1.07 kcal/mol, corresponding to weaker observed binding for PUM1 than PUM2; the RMSE indicates the error after subtracting the constant offset. (B) Predicting PUM1 binding with the PUM2-based model. Inset shows the distribution of deviations from predicted values. (C) Schematic representation of the single amino-acid change in PUM1 repeat ‘R6’. (D) Differences between the single-mutant specificities of wild-type and mutant PUM1. Differences of median single mutant penalties are shown, and the error bars indicate propagated 95% CIs. “N.A.” indicates lack of detectable binding. (E) Predicted mutant PUM1 affinities (based on the PUM2 model) versus observed affinities; the ΔΔG values are relative to the UGUAUAUAU consensus. (F) Predicted versus observed mutant PUM1 affinities with the altered penalty for 6U (ΔΔG = –0.35 kcal/mol instead of +1.49 kcal/mol). See also Figure S6.

### Evaluating the precision of Pumilio engineering

The modular structure of PUF proteins has made them attractive platforms for engineering new RNA specificities (Abil et al., 2014; Adamala et al., 2016; Campbell et al., 2012; Campbell et al., 2014; Cheong and Hall, 2006; Dong et al., 2011; Filipovska et al., 2011; Porter et al., 2015; Wang et al., 2002; Weidmann and Goldstrohm, 2012; Zhao et al., 2018). Given our observation of complexity of the PUF protein specificity landscape not captured by a simple linear motif and the low-throughout nature of most previous studies, we aimed to comprehensively evaluate the precision of PUF protein engineering using a previously designed PUM1 mutant (Cheong and Hall, 2006).

We investigated the specificity of a PUM1 mutant designed to alter specificity for position 6 in the RNA from A to U through a single amino acid substitution in Repeat ‘6’ (Cheong and Hall, 2006). The C935N mutation changes the characteristic A-specific triad of amino acids (QRxx**C)** to resemble the conserved residues of a U-specific repeat (QYxx**N**; Figure 5C), and this mutant was previously shown to have 6-fold greater affinity for the UGUAU**U**UA sequence than for the wild-type UGUAU**A**UA consensus (Cheong and Hall, 2006). Analysis of single mutant penalties relative to wild-type PUM1 confirmed a change in mutant PUM1 specificity at position 6: while wild-type PUM1 bound the 6U RNA 12-fold weaker than it bound the 6A consensus, the engineered protein showed a 1.8-fold preference for 6U instead, giving a 22-fold specificity change (Figure 5D). The relative affinities were unaffected for the other RNA residues at position 6 (6C, 6G), and no other single mutants showed a significant difference, supporting a local effect from the PUM1 mutation (Figure 5D).

If the programmed affinity change is fully local, then our model, based on wild-type PUM1/2, should accurately predict binding, except for sequences containing a U at bound position 6. Applying the original PUM1/2 model to mutant PUM1 binding led to accurate predictions for most variants, but 18% of variants bound considerably (>1 kcal/mol) more tightly than predicted (Figure 5E). Changing a single term in our binding model to account for the altered 6U penalty for mutant PUM1 (–0.35 kcal/mol instead of +1.49 kcal/mol) gave accurate predictions for these outliers (Figure 5F), consistent with a localized effect of the PUM1 mutation on binding of RNA variants that contain U at position 6. The modified thermodynamic model predicted affinities for 99% of variants within 1 kcal/mol of their observed binding free energy and had an RMSE of 0.32 kcal/mol, similar to that for wild-type PUM1 (Figures 5F,B). Thus, our quantitative model can be readily modified and applied to new PUF proteins.

### Assessing the thermodynamic model for PUM2 RNA occupancies in vivo

The occupancy of PUM2 at RNA sites in vivo could simply reflect the relative thermodynamic affinity for sites within each RNA, and because of the large excess of consensus RNA binding sites over PUM1/2 proteins in human cells (Lee et al., 2016), a strictly thermodynamic binding model would predict a linear relationship between site affinity and PUM2 occupancy. Alternatively, if RNA chaperones or helicases were to displace PUM2 faster than its intrinsic dissociation rate, binding sites with different affinities but the same association rate constants would have the same occupancies (Figure S7A,B). To assess the extent to which in vivo binding is driven thermodynamically, we compared predictions from our thermodynamic binding model to published in vivo UV crosslinking measurements for PUM2 from the ENCODE project (Consortium, 2012; Van Nostrand et al., 2016).

We evaluated PUM2 binding data, obtained by enhanced UV crosslinking and immunoprecipitation (eCLIP), in bins of predicted affinity, because quantification of individual RNA sites is currently limited by low sequencing depths and may be further subject to experimental biases (Darnell, 2010; Sugimoto et al., 2012; Wheeler et al., 2018). Putative PUM2 binding sites within expressed mRNAs were identified as sites with predicted binding affinities within 4.0 kcal/mol (∼1000-fold) of the consensus sequence (Methods). eCLIP signal was summed across an 80nt window surrounding each site and divided by the relative expression of its transcript. Strikingly, we observed quantitative agreement between relative affinities predicted by our thermodynamic model and the median eCLIP enrichment signal across the predicted affinity bins (Figure 6A, points vs. dashed line). Close agreement was observed for predicted sites both with and without flipped residues (Figure S7D). Thus, the in vivo binding data are consistent with thermodynamically driven occupancy, and the binding sequences and modes identified by RNA-MaP are bound in vivo at the levels expected based on their affinities.

**Figure 6.**
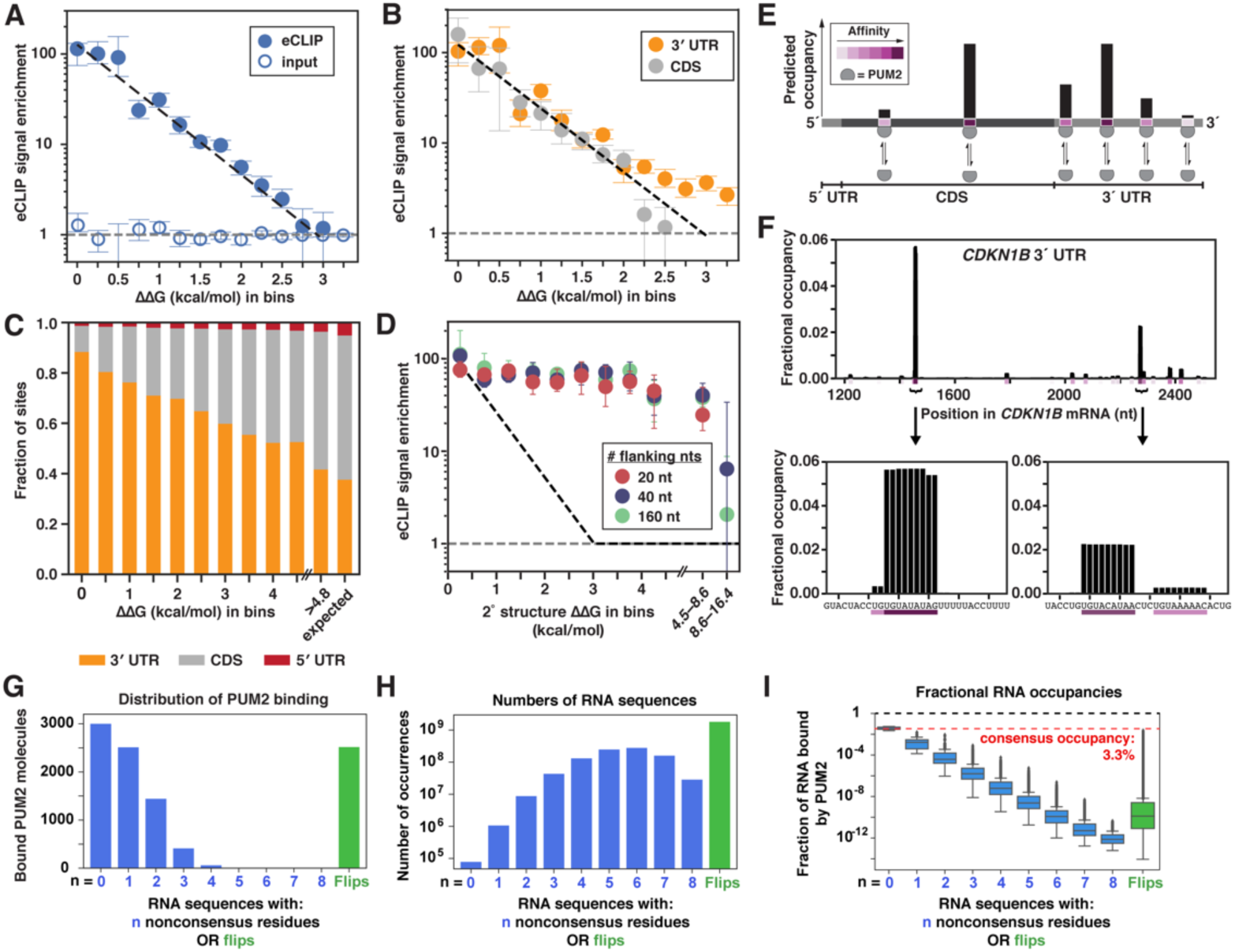
Testing the Thermodynamic Model in Vivo.

(A) Scatterplot comparing thermodynamic affinity predictions with eCLIP enrichment in K562 cells (Van Nostrand et al., 2016). Median eCLIP enrichments across sites within bins of predicted relative affinities are shown, and error bars indicate 95% CIs on the median. Only sites lacking adjacent UGUA-containing sites (within 100 nt) are shown, due to inflation of eCLIP signal observed in the presence of nearby sites (Figure S7C). Black dashed line indicates the predicted change in eCLIP signal with increasing predicted ΔΔG values, relative to the eCLIP signal in the lowest ΔΔG bin. eCLIP (closed circles) and input (open circles) correspond to crosslinked samples that were or were not treated with anti-PUM2 antibody, respectively (Van Nostrand et al., 2016). The grey dashed line indicates the eCLIP enrichment for sites with predicted ΔΔG values greater than 4.5 kcal/mol (expressed transcripts); since eCLIP signal and input were each normalized to this value, this expected enrichment is equal to 1.

(B) Comparing eCLIP enrichment for sites within 3´ UTR (orange) or CDS (grey) of expressed genes in K562 cells. Only sites that are at least 100nt from the nearest UGUA are included. Black and grey lines are as in *A*.

(C) Fractions of sites within bins of predicted ΔΔG values that are annotated as 3´ UTR, CDS, or 5´ UTR.

(D) Comparing the median eCLIP enrichment of consensus sites in bins of predicted secondary structure stabilities for structures blocking the PUM2 consensus site (Figure 2C; Methods). Colors indicate the number of flanking nucleotides (nt) included in the stability calculations. Dashed line indicates the predicted change in eCLIP signal for increasing secondary structure stability. Medians and 95% CIs for bins with at least 20 sites are shown.

(E) Predicting PUM1/2 binding occupancies of cellular RNAs. Schematic representation of the output of an occupancy prediction algorithm.

(F) Example of thermodynamic occupancy predictions for the 3´ UTR region of the human cyclin dependent kinase inhibitor 1b *CDKN1B* mRNA, a known PUM1/2 target (Kedde et al., 2010). The fractional occupancies on the Y axis account for cellular PUM2 and RNA abundances (Methods).

(G) PUM2 binding landscape across the human transcriptome, predicted by our thermodynamic model using in vivo PUM2 and mRNA levels (see Methods). Bars indicate the number of bound PUM2 molecules across RNA binding sites with 0–8 nonconsensus residues without flipped residues (blue) or with up to two flipped residues (green). The consensus was defined as UGUA[ACU]AUAN.

(H) Total counts of 9–11mers with the indicated numbers of nonconsensus residues and/or flips in the human protein-coding transcriptome.

(I) The fraction of each RNA site (from part *A*) that is bound by PUM2. Boxes indicate the interquartile range, and the spread arises because different nonconsensus residues give different amounts of destabilization.

See also Figure S7.

Because Pumilio proteins have generally been identified to act via 3´ untranslated regions (UTRs), we wondered whether there might be lower average occupancy in coding sequences (CDS) and in 5´ UTRs. One of several possible factors that could lead to such disparities would be PUM2 dissociation stimulated by ribosomes translating through these binding sites, as this would decrease CDS but not 3´ UTR occupancies. Nevertheless, comparison of the occupancy around PUM2 sites in 3´ UTRs and CDSs showed indistinguishable eCLIP enrichments (Figure 6B; 5′–UTR sites were not included because of the small number of predicted sites in this region). This comparison suggests that the inherent thermodynamic stability of a site is the overarching driver of in vivo occupancy, rather than the location of the site within the mRNA.

We observed a strong enrichment of PUM2 consensus sites in 3´ UTR sequences relative to CDS and the 5´ UTR, with ∼90% of the strongest binding sites (ΔΔG < 0.5 kcal/mol) located within 3´ UTRs, despite 3′ UTRs constituting on average only 35% of the mRNA length (Figure 6C, Figure S7E (Consortium, 2012)). Enrichment was evident, though diminished, for sites with weaker predicted affinities, for ΔΔG values up to ∼3 kcal/mol. This enrichment is consistent with the established functional roles for PUM2 via 3´ UTR binding ((Bohn et al., 2018; Kedde et al., 2010; Miles et al., 2012; Rodrigues et al., 2016; Wickens et al., 2002)).

Our in vitro thermodynamic measurements indicated that RNA secondary structure formation can strongly limit the accessibility of PUM2 binding sites, and thus decreases PUM2 binding (Figure 2C,D; I.J., W.R.B. et al., in prep.). In contrast, when we compared the in vivo eCLIP signal around consensus sites with varying predicted structure content, the eCLIP signal was largely unaffected, with no change in occupancy for predicted structural stability of up to ∼4 kcal/mol (Figure 6D; 37 °C). This result was independent of the number of flanking nucleotides included in the structure prediction, from 20 nt to 160 nt. A rare subset (<2%) of sites had very high predicted structure stability (ΔΔG_fold_ > 8.6 kcal/mol; 160 nt window) and showed slightly diminished eCLIP signal, suggesting that highly stable structures may lead to decreased binding. However, RNA structure effects on the vast majority of PUM2 sites appear to be negligible in vivo.

## Discussion

RNA/protein interactions are integral to regulation of gene expression (Singh et al., 2015). To define and predict the complex networks of RNA/protein interactions, quantitative descriptions of RNA/RBP thermodynamics are needed. As an early but critical step toward this goal, we have built a predictive model for RNA binding by the important human RBPs PUM1 and PUM2. Our direct thermodynamic binding measurements for thousands of RNAs and the corresponding model provide testable predictions of in vivo RNA interactions, a platform for analyzing specificity of related RBPs, and novel biological and biophysical insights.

### Applications to cellular interactions and RNA properties

Our thermodynamic model provides a quantitative foundation for testing cellular factors that influence RNA behavior, as they can be read out as deviations from the thermodynamically predicted behavior.

Comparison of predictions from our binding model to published in vivo cross-linking data (Van Nostrand et al., 2016) supports the simple notion that thermodynamics is a prime driver in determining RNA occupancy for PUM2. While it would be surprising if thermodynamic affinities did not, at some level, influence RNA binding in vivo, other models are possible. For example, rapid Pumilio protein dissociation by the action of RNA helicases would level occupancies for all sites above a certain threshold affinity (Figure S7B), and translation by ribosomes that displaces PUM2 proteins faster than equilibration of PUM2 binding would yield CDS occupancies lower than 3´ UTR occupancies. However, a close correspondence between thermodynamic predictions and extents of crosslinking in vivo provides evidence against these alternate models (Figure 6A, B). Ribosomes traverse CDS sites, and presumably displace bound factors, about once every 10 seconds (∼0.1 s^−1^ (Halstead et al., 2015; Schwanhausser et al., 2013)). This rate sets an upper limit for the equilibration time for PUM2 binding—it must occur faster than ribosomal displacement for us to observe no difference in occupancy in CDSs compared to 3´ UTRs. This rough lower limit estimate is similar to the rate constant for dissociation of PUM2 from the consensus sequence in vitro (∼0.1 s^−1^; 37 °C; unpublished results). This calculation suggests that PUM2 dissociates from consensus sites on the timescale of translation, and faster for nonconsensus sites, in the absence of ribosome displacement or other stimulating factors, consistent with our observations. More direct assessments of the interplay between thermodynamic and dynamic factors in vivo will be of great interest, given the potential for transcript-specific effects and the need to determine whether cellular conditions and factors alter binding and dissociation rates.

PUM2 binding to ssRNA and the expectation, supported by in vitro data, that RNA structure inhibits binding (Figure 2C, D; I.J., W.R.B. et al., in prep.) provided us with an opportunity to assess the stability of predicted RNA structures in cells. Recent studies have suggested that RNA structure is destabilized in the cellular milieu (Ding et al., 2014; Guo and Bartel, 2016; Rouskin et al., 2014; Spitale et al., 2015), but given the fundamental importance of determining the physical state and dynamic behavior of molecules in cells, additional approaches and data to address this question are called for. Using published eCLIP data (Van Nostrand et al., 2016), we observed that binding sites predicted to be minimally accessible (<0.1%; ΔΔG_fold_ ≳ 4 kcal/mol) gave enrichments that were essentially indistinguishable from sites predicted to lack structure (Figure 6D). These results provide independent support for structure disruption in vivo, which could result from a high density of bound RBPs that outcompetes RNA structure formation and/or the action of RNA chaperones. Moving forward, coupling of in vitro thermodynamic measurements with quantitative in vivo analysis will aid in determination of cellular factors responsible for destabilizing RNA structure and the degree to which structure varies across RNAs.

While the eCLIP data (Van Nostrand et al., 2016) enabled us to perform initial tests of thermodynamic predictions for PUM2 in vivo, future tests will be needed to assess factors that lead to occupancy variation at the level of individual RNAs and to confidently distinguish variation caused by biological factors vs. technical artifacts (Wheeler et al., 2018). Improvements in individual-RNA signal intensity and quantitative controls will allow dissection of these factors.

### An algorithm for predicting Pumilio occupancies

Our comprehensive thermodynamic PUM2 binding model, along with estimated in vivo PUM2 and RNA levels, allows prediction of PUM2 occupancy across the entire transcriptome. We supply a computational algorithm to carry out these predictions, and depict the output of this tool for one transcript in Figure 6E,F. Our algorithm can be used to predict occupancies for individual sites and to design tests for affinity, local structure, and other factors that might affect RBP binding in vivo. The model can be extended to predict global changes in occupancies in response to changes in protein levels, affinities, or specificities.

Our PUM2 binding model was readily extended to PUM1, which has identical specificity, and readily adapted for an engineered PUM1 variant, by adjusting a single term to reflect the changed specificity (Figures 5C–F; (Cheong and Hall, 2006)). Similarly, the model, with additional terms and adjustments, should account for specificity of the spectrum of Pumilio proteins, and the approach executed herein should be extensible to provide quantitative affinity for many additional RBPs.

### Specificity and cellular RNA/protein binding landscapes

Determining quantitative RBP binding landscapes and how these landscapes change with changes in RBP and RNA levels, modifications, and other cellular factors is a critical component of a complete description of the RBP/RNA interactome. The shape of the binding landscape––i.e., RBP occupancies across RNA sequences present in a cell––has implications for regulation, evolution, and engineering. We illustrate some of these implications, including non-obvious consequences of binding specificity.

Figure 6G shows predicted PUM2 binding across all possible RNA binding sites in the human protein-coding transcriptome (Methods); RNA sequences are defined by the number of nonconsensus residues relative to the UGUA[ACU]AUAN consensus and by the presence or absence of flipped residues. Despite the stronger binding of PUM2 to consensus than other sequences, our thermodynamic model predicts that less than a third of cellular PUM2 is bound to consensus sites (Figure 6G; Methods). Instead, the majority of the protein is distributed across the much more common nonconsensus sites that contain up to 4 nonconsensus residues and/or flipped residues that still have moderate affinities (Figure 6G,H). The varied, moderate, nonconsensus residue penalties of PUM2 allow for a smooth gradient of binding occupancies (Figure 6I; see also Figure 3G and Figure 6A; Table 1). Thus, we speculate that regulatory response strengths to PUM2 (e.g., in RNA decay) can be finely tuned at the level of RBP binding, by utilizing different RNA sequences along a gradient of affinities.

A less obvious influence of RBP specificity on regulation arises because the binding to nonconsensus RNA sites reduces the pool of protein available to bind to consensus sites. Because binding occurs across a large number of sites, and because of the much greater number of potential binding sites than the number of PUM2 molecules in cells, only a small fraction of each consensus site (UGUA[ACU]AUAN) is predicted to be occupied by protein (∼3%; Figure 6I). Thus, even in the presence of *total* protein concentration in excess of the consensus affinity ([PUM2] = 10 nM vs. *K*_D_ = 3 nM at 37 °C), binding is decidedly subsaturating. This subsaturating binding renders per-site occupancies highly sensitive to changes in PUM2 levels or affinity, analogously to cellular enzymes that operate in a subsaturating regime ([Substrate] ∼ *K*_M_) to enable greater sensitivity to cellular changes in substrate concentration (Berg JM, 2002). In contrast, in a hypothetical extreme scenario in which PUM2 bound only a single 9mer consensus sequence with infinite specificity (UGUAUAUAG; n ≈ 4000), these consensus sites would remain fully occupied across a range of PUM2 concentrations (with PUM2 in excess of these sites), making them effectively insensitive to changes in total or active PUM2.

Additional advantages from a near-continuous landscape of occupancies may arise because natural selection can ‘choose’ between sequences with very similar binding to co-optimize for other properties, such as binding to other RBPs, RNA structure, and tuning of translation. Also, by having different sequences with similar affinities, mRNA targets could be sub-divided into groups that are subject to distinct control (e.g., by covalent modification of one but not the other RNA sequence), thereby creating a more precise and interconnected regulatory network.

Finally, the near-continuous landscape of PUM2 occupancies across mRNA sequences would be expected to render it highly evolvable. The presence of a large number of sites bound with moderate affinities, and the often subtle effects of individual substitutions should allow evolution to tune regulation of existing sites as RNA sequences undergo mutation and will increase the ability to co-opt new RNAs for regulation. Indeed, PUM2 orthologs are present throughout Eukaryotes that recognize different sets of RNAs; these transitions that occurred multiple times in evolution may have been facilitated by PUM2’s moderate specificity (Gerber et al., 2006; Hogan et al., 2015; Jiang et al., 2010).

### Biophysical insights into Pumilio•RNA interactions

Quantitative high-throughput thermodynamic data can also help the development of testable physical models. Our data revealed that the A-recognition modules at positions 4, 6, and 8 give highly similar specificities, whereas the U-recognition modules at positions 1, 3, 5, and 7 vary dramatically in their specificities, from no discrimination against A and C at position 5 to ∼10^2^ fold specificity at positions 1 and 3 (Figure 2E, Figure S4H–J). The differential specificity across the U-recognition modules could arise, at least in part, from differences in orientations and constraints imposed by the different neighboring positions, which can allow more or less optimal positioning at each U-recognition module. Similarly, the slightly weaker discrimination at position ‘8’, at the end of the Pumilio modules, relative to the internal A-recognition modules, may arise because there are fewer conformational restraints 3´ of this position, allowing noncognate bases to more readily find alternative bound conformations. These observations are of practical importance for engineering Pumilio proteins and of broader importance for understanding and modeling RNA recognition by proteins.

The absence of measurable coupling between most neighboring residues suggests that the orientation of entry of the RNA into a site is not generally affected by the identity of the neighboring residue. The simplest model is that cognate and noncognate residues are bound with similar backbone configurations (or ranges of backbone conformations), and this interpretation is consistent with crystallographic observations that both cognate and noncognate bases dock into Pumilio binding sites and that backbone trajectories leading into and out of Pumilio repeat sites are similar with cognate and noncognate bases (Gupta et al., 2008; Lu and Hall, 2011; Wang et al., 2009). More generally, the observed energetic independence between most adjacent RNA residues suggests that there is sufficient room in the binding sites and/or sufficient degrees of freedom in the RNA backbone to allow the backbone to ‘forget’ its specific interactions at the adjacent sites. Nevertheless, a subset of positions give coupling, and coupling is likely more prevalent for at least a subset of other RBPs.

Whereas neighboring bound residues largely do not affect each other, larger energetic effects are observed in cases of inserted residues that can flip away from the recognition sites, (Figure 3E and Table 1). A residue that follows a flipped residue will experience a larger loss in conformational entropy upon docking than a residue that follows and is positionally restricted by a preceding docked residue. Nevertheless, the specificity for neighbors is the same whether or not there is an intervening flipped residue—i.e., the same free energy terms can be used for each bound residue (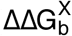, Table 1A) whether or not there is a flipped residue and associated flipping penalty (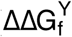; Table 1B). The observed energetic constancy suggests that flipped residues do not significantly alter the docked states for neighboring cognate and noncognate residues and that there are no alternative bound states for the neighboring residues that are both reachable and more favorable energetically than the standard docked state.

These brief considerations of the interrelationship between binding thermodynamics and conformational properties underscore complexities in understanding and modeling RNA/RBP interactions. Relative to the conformationally more constrained dsDNA and studies of DNA/protein interactions, broader ensembles are expected for ssRNA, both free and bound to RBPs, highlighting the enormous challenge faced in modeling RNA/RBP binding affinities and specificities. Considering the large number of proteins that bind RNA, the ability to ultimately define binding landscapes for all RBPs will likely require computational approaches, and developing models that can accurately predict thermodynamics we believe will require guidance and testing with large accurate thermodynamic datasets, such as those obtained herein.

## Acknowledgements

We thank Wipapat Kladwang for outstanding technical assistance with emulsion PCR. We thank Namita Bisaria, Greg Hogan, Julia Salzman, Erik Van Nostrand and members of the Herschlag lab for helpful discussions and comments on the manuscript. This work was funded by grants from the U.S. National Institutes of Health: P01 GM066275 (D.H., R.D., W.J.G.), R35 GM122579 (R.D.), R01 GM121487 and R01 GM111990 to (W.J.G), and by the Beckman Center. W.J.G acknowledges support as a Chan-Zuckerberg Investigator.

## Author contributions

Conceptualization: IJ, PPV, SKD, WRB, KK, DH, WJG; performing experiments: PPV, IJ, RS; formal analysis: SKD, WRB, PPV, IJ, KK, VS; resources (imaging station design, maintenance and analysis tools): JOA, CJL; writing—original draft: IJ, SKD, WRB, PPV, DH, WJG; writing—final draft: IJ, SKD, WRB, PPV, KK, JOA, CJL, VS, RS, RD, DH, WJG; project administration: DH, WJG; funding acquisition: DH, WJG, RD.

## Supplemental Figures

**Figure S1.**
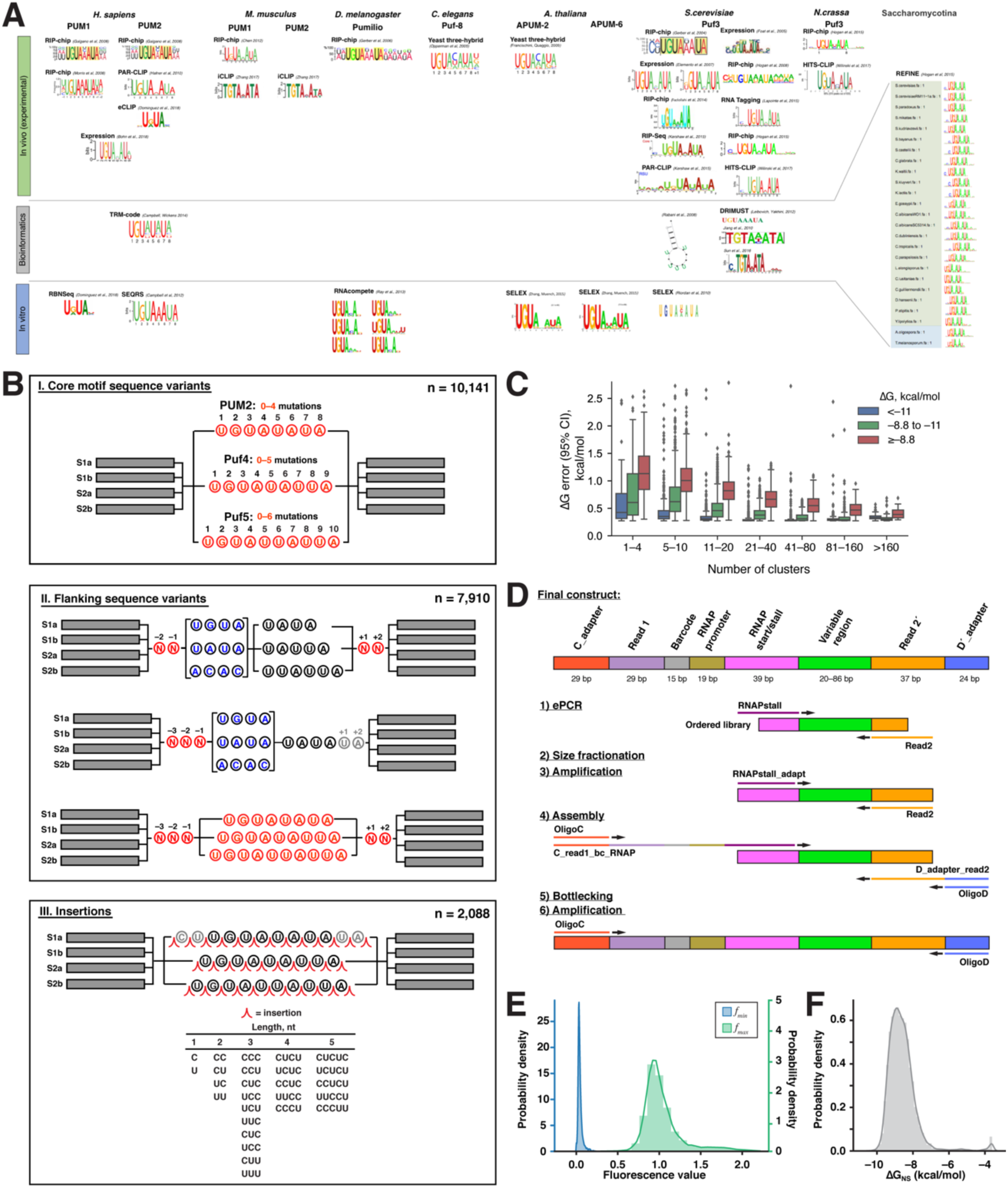
RNA Array Library Design, Preparation and Experiments, Related to Figure 1 and Methods. (A) Previously identified sequence motifs of human PUM1, PUM2 and orthologous PUF proteins (Bohn et al., 2018; Campbell et al., 2012; Campbell et al., 2014; Chen et al., 2012; Dominguez et al., 2018; Elemento et al., 2007; Fazlollahi et al., 2014; Foat et al., 2005; Francischini and Quaggio, 2009; Galgano et al., 2008; Gerber et al., 2004; Gerber et al., 2006; Hafner et al., 2010; Hogan et al., 2015; Jiang et al., 2010; Kershaw et al., 2015; Klass, 2013; Lapointe et al., 2015; Leibovich and Yakhini, 2012; Morris et al., 2008; Opperman et al., 2005; Rabani et al., 2008; Ray et al., 2013; Riordan et al., 2011; Sun et al., 2016; Wilinski et al., 2017; Zhang and Muench, 2015; Zhang et al., 2017). (B) RNA libraries used in this study. Three broad categories of PUM2 binding site variants were designed: (I) mutations in the core binding site; (II) flanking sequence variants; (III) insertions. Each variant was represented in 2–4 scaffolds (S1a–S2b; Figure 1B). Within each category, the design was based on the previously established consensus motif of PUM2 and its orthologs; substantial additional sequence variation was achieved by applying the same types of perturbations to the consensus motifs of related PUF proteins, *S. cerevisiae* Puf4 and Puf5 ((Gerber et al., 2004; Hogan et al., 2008; Hogan et al., 2015; Riordan et al., 2011; Wilinski et al., 2015)). To prevent a preponderance of weak binders, in the mutation library (I) no more than two mutations were allowed in the highly conserved UGUA core of the PUF motifs, with the exception of negative controls in which UGUA was mutated to ACAC. Numbers of designed variants in each category are indicated. (C) Affinity and number of clusters per variant affect measurement certainty. Box plots show measurement uncertainties from PUM2 replicate 1, with the box indicating the interquartile range, the whiskers indicating the minimum and maximum values and the points indicating outliers. Only variants with ≥5 clusters and ΔG values below –8.8 kcal/mol were included in subsequent analyses (see Methods). (D) Preparation of DNA array library. See Methods for detailed sequence information and description of individual steps. (E, F) Distribution of initial fit parameters for variant-independent parameters *f*_min_, *f*_max_, and ΔG_NS_. (E) Histogram of the initial per-variant values for *f*_min_ (blue) and *f*_max_ (green) for variants that reached near-saturation of binding (>95% bound at highest concentration based on initial per-variant *K*_D_) and that met other criteria for high-confidence binders during initial fitting, as defined in Methods. (F) Histogram of the initial per-variant values for ΔG_NS_ (= RT ln*K*_D,NS_; Methods). Values for PUM2 replicate 1 are shown.

**Figure S2.**
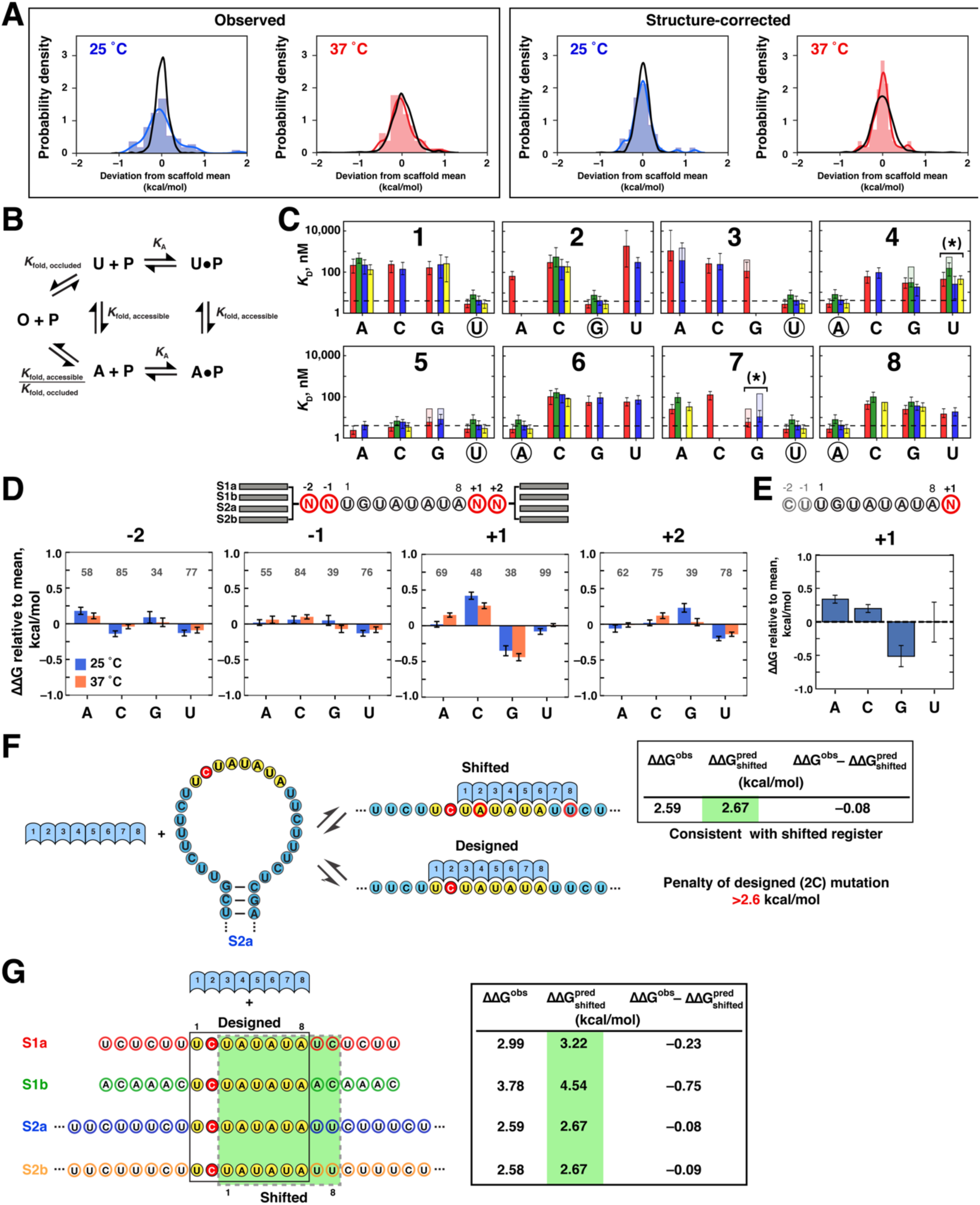
Analysis of Single-Mutant Variant Binding to PUM2, Related to Figure 2. (A) Assessing significance of scaffold differences. For each binding site variant (single mutants and UGUAUAUA consensus), we determined the deviations of individual scaffold affinities from the scaffold mean for that variant. The distribution of these deviations was then compared to the distribution expected from experimental error (black line). The panels show distributions at 25 °C (blue) and 37 °C (red), before (left) and after (right) accounting for structure effects, as predicted by RNAfold ((Lorenz et al., 2011); see Methods). A summary of the standard deviations of the displayed distributions is shown in Figure 2B. (B) Model for accounting for RNA structure effects on PUM2 binding. U: unfolded RNA; P: protein; A: accessible RNA (i.e., structured RNA with the PUM2 binding site accessible); O: occluded RNA (i.e., PUM2 binding site occluded by base-pairing). A and O represent ensembles of the respective states; see Figure 2C). *K*_A_: PUM2 association constant for the unstructured binding site (assumed to be the same for binding to U and A, but no protein binding occurs to O); *K*_fold,accessible_: equilibrium constant for RNA folding from the unfolded state to the ensemble of structures in which the PUM2 binding site is accessible (determined by constraining the binding site to the single-stranded state in Vienna RNAfold; 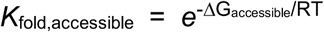); *K*_fold,occluded_: equilibrium constant for RNA folding from the unfolded state to the ensemble of structures in which the PUM2 binding site is occluded by structure. (C) PUM2 affinities for UGUAUAUA single mutants at 37 °C. Solid bars indicate mutant affinities after accounting for structure effects, and transparent regions indicate predicted structure effects. Asterisks indicate mutants for which affinities measured in different scaffolds were significantly different (10% FDR) prior to accounting for structure effects (differences between total bar heights). No significant differences remained after accounting for structure effects using RNAfold. (D) Flanking sequence effects on PUM2 binding. *Top*: Library design. Two bases upstream and downstream from the UGUAUAUA consensus were randomized (N=A/C/G/U; n = 256) and the resulting sequences were embedded in 2–4 scaffolds each. *Bottom*: Effects of each base at the indicated positions relative to the average affinity. The effects were determined by calculating the average affinity of variants containing the indicated base (regardless of the identities of other flanking bases) and subtracting the mean of all four bases. Error bars indicate 95% CI of the mean. Only variants with predicted structural effects <–0.5 kcal/mol were included, and the numbers of variants containing each base are indicated above the bars. Blue and orange bars indicate the sequence effects at 25 °C and 37 °C, respectively. The observed sequence effects at positions −2, −1 and +2 are small and are further dampened at 37 °C, suggesting that residual structural effects are responsible for the small observed effect. The strongest effect was observed at position +1, and the observed preference for G was maintained at 37 °C, indicating a weak, base-specific interaction. (E) Gel-shift measurements of sequence effects at position +1. *Top*: Oligonucleotide design; N = A/C/G/U. *Bottom*: Effects of each base relative to the mean affinity for all four variants. Averages and standard errors from two replicate experiments are shown. (F) Assessment of alternative binding registers. Example of preferential binding in an alternative register accompanying a destabilizing mutation in the designed binding site. Affinity effects are shown as ΔΔG values, relative to the average affinity for the UGUAUAUA consensus across scaffolds. The predicted ΔΔG for the alternative register corresponds to the sum of the ΔΔG values for the two nonconsensus residues, 2A and 8U (red outlines), assuming energetic independence. Given that the observed ΔΔG (ΔΔG_obs_) matches the prediction for the alternative register, the measurement only establishes a lower limit for the penalty of the intended, 2C, mutation. (G) Observed and predicted ΔΔG values for the best alternative register 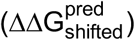 in each of the four scaffolds for the 2C variant (other variants with predicted alternative registers are listed in Table S1). The close agreement of alternative register predictions with observed affinities in the S1a, S2a, and S2b scaffolds suggests predominant binding in the shifted register, whereas the 0.75 kcal/mol difference in the S1b scaffold suggests preferential binding in the intended register, such that the observed value of 3.78 kcal/mol reflects the real effect of the 2C mutation.

**Figure S3.**
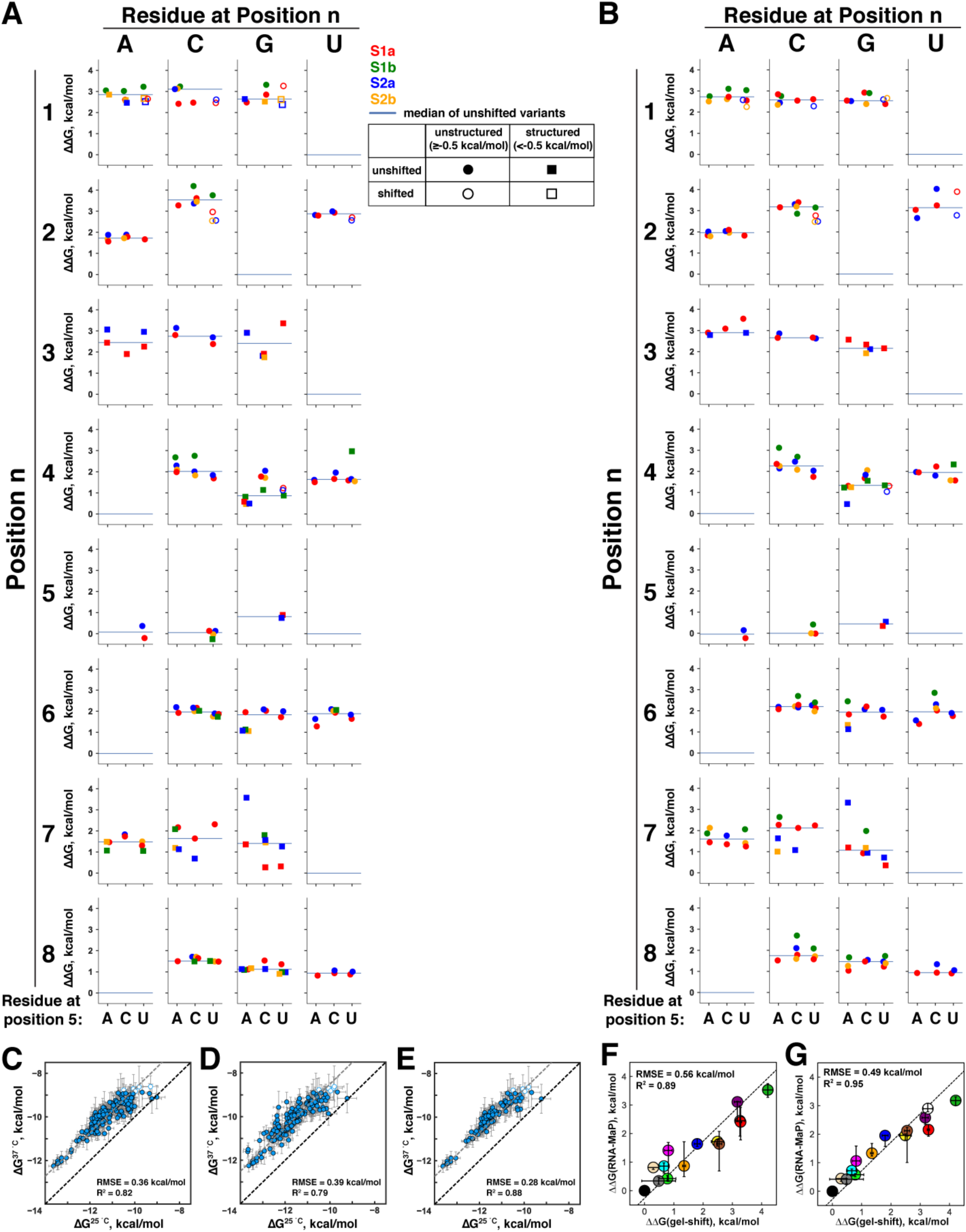
Determination of Single Mutant Penalties, Related to Figure 2. (A, B) Single mutation effects on PUM2 binding across backgrounds of different position 5 identities (5 = A/C/U) at 25 °C (A) and 37 °C (B). Symbols indicate the ΔΔG values for each mutation relative to the weighted mean of UGUA[A/C/U]AUA affinities across scaffolds (color-coded). As shown in Figure 2D and previously (Lu and Hall, 2011), residues A, C and U are bound with identical affinities at position 5, and other positions show high consistency of mutational penalties measured in 5A/C/U backgrounds, consistent with energetic additivity with respect to position 5. Thus, mutational penalties from all three backgrounds were used for determination of single mutant penalties, with median ΔΔG values indicated by horizontal lines. Circles indicate unstructured variants, while squares denote variants with predicted structure effects greater than 0.5 kcal/mol, and the displayed values correspond to structure-corrected affinities. Open symbols indicate variants that have predicted alternative registers within 1 kcal/mol from the observed ΔΔG value; these variants were not included in determining the median. None of the variants in the 5A/C backgrounds were predicted to have alternative binding registers. (C–E) Scatterplots of RNA single mutant affinities for PUM2 at 25 °C and 37 °C. (C) Observed ΔG values for single mutants in position 5A/C/U backgrounds (n = 161). Open symbols indicate variants with affinities outside the reliable affinity threshold (−8.8 kcal/mol; not included in RMSE and R^2^ determination). The grey dashed line indicates the mean offset between values measured at 25 °C and 37 °C, corresponding to a (20±10)-fold decrease in affinity at 37 °C. (D) ΔG values after accounting for predicted structure (RNAfold). The poorer correlation than observed in part *C* is likely due to limited accuracy of structure predictions (I.J., W.R.B. et al., in prep). (E) Correlation of observed ΔG values at 25 °C and 37 °C for variants with no predicted structure (ΔG_fold_ > –0.5 kcal/mol; n = 107). The correlation approaches that of replicate measurements (Figure 1E; RMSE = 0.26 kcal/mol; R^2^ = 0.96). (F, G) Comparison of single mutant affinities measured by RNA-MaP and gel-shift (25 °C). 1C: purple, 2A: yellow, 2C: green, 3A: white, 3G: red, 4G: orange, 4U: blue, 5G: wheat, 7C: brown, 7G: magenta, 9A: lime, 9C: cyan, 9U: grey. RNA-MaP values are medians and 95% CIs of individual mutation penalties in 5A/C/U backgrounds at 25 °C (F) and 37 °C (G) (Figure S3A,B and Figure 2E; the RNA-MaP medians were determined after accounting for secondary structure and shifting). The gel-shift values for RNAs with mutations in positions 1–8 were determined by competition measurements of 8mer oligonucleotides, with mutations introduced in the UGUAUAUA background, and the ΔΔG values were calculated relative to this UGUAUAUA consensus. Position 9 mutants are based on direct measurements of CUUGUAUAUAN oligonucleotides, relative to the most tightly bound residue (G; Figure S2D). The better agreement of gel-shift values with 37 °C RNA-MaP values is consistent with small structural effects on RNA-MaP-derived values.

**Figure S4.**
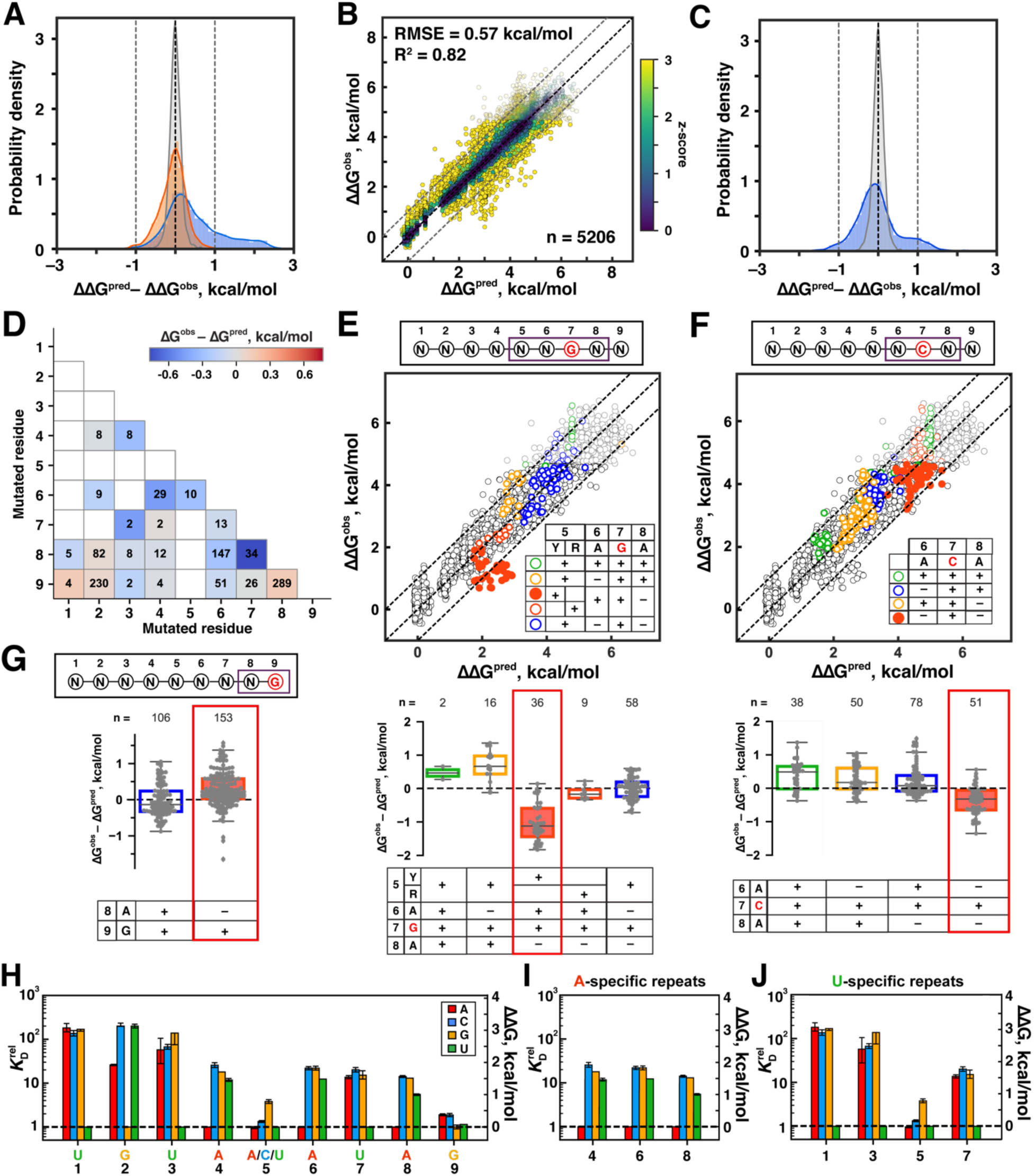
Developing a Predictive Model for PUM2 Specificity, Related to Figure 3. (A) Distribution of differences between model predictions and observed affinities through PUM2-binding model development. Blue: additive consecutive model (Figure 3A); orange: final model including flipping and coupling terms (Figure 3G); grey: the distribution of standard errors of measured ΔΔG values. Coupling did not noticeably affect the distribution, so the additive nonconsecutive model without coupling (Figure 3F) is not displayed. (B) Global fit to the additive consecutive model. Parameter constraints were identical to those used for fitting the additive nonconsecutive model (Figure 3F) and corresponded to the 95% confidence intervals from single mutant measurements (Figure 2E) or ±0.4 kcal/mol, whichever was greater. Scatter plot of the predicted versus observed ΔΔG values is shown. The points are colored based on their z-scores for the deviation from predicted value (see Figure 3A). The black dashed line indicates perfect agreement with prediction, and dashed grey lines denote 1 kcal/mol deviation from predictions. While the additive consecutive model performed considerably better with globally fit parameters than with experimentally determined single mutant penalties (Figure 3A), there was substantial asymmetry and ten percent of variants had predicted affinities >1 kcal/mol from the observed value (versus 0.00% expected from measurement error); see part C. (C) Blue: distribution of differences between values predicted by the additive consecutive model (using parameter values determined by global fit; part B) vs. observed values; grey: distribution of standard errors of measured affinities, as in part A. (D) Double mutant coupling at 25 °C. Colors indicate the difference of observed ΔΔG values (ΔΔG_obs_) from the sum of ΔΔG values for the individual mutations (ΔΔG_pred_ = ΔΔG_1_ + ΔΔG_2_). Double mutants were defined as all sequences differing from the UGUA[A/C/U]AUAU reference sequence at two positions and predicted to be in the most favorable register with no flips (Methods); only variants with predicted structure effects of <0.5 kcal/mol were included. Numbers of variants containing each combination of mutations are shown inside the respective boxes; only mutants represented by more than one variant in our library are shown. White boxes indicate absence of the respective mutants in the best register in our unstructured library: these absences are primarily due to the highly destabilizing effects of mutations in the 5′ half of the motif, which in combination with a second mutation lead to binding outside our reliably measurable range and/or to shifting of the preferred binding site to an alternative binding register; low representation of 5G-containing mutants is due to predicted structure. (E) Coupling between the 7G mutation and adjacent positions, assessed as the extent to which affinities for variants with the indicated residue combinations (inset) deviate from predicted values. The predictions are based on the additive nonconsecutive model (Figure 3F). All sequences containing the indicated combination of residues (inset) in the predicted best consecutive register were considered. Black residues in the inset table indicate consensus residues. The top panel shows the variants overlaid on the scatterplot of predicted and observed values for the entire library (same as Figure 3F). The bottom panel shows the distributions of differences from predicted values for each group of variants. The coupling term that was incorporated in the final model is highlighted by solid red circles (top) and red outline (bottom). To avoid overfitting, only most pronounced instances of apparent coupling were included in the final model. (F) Coupling between the 7C mutation and flanking residues, as in *E*. (G) Coupling between positions 8 and 9. As in *E* and *F*, coupling was assessed as the extent to which affinities for variants with the indicated residues deviated from predicted values. The panel illustrates the loss of the small stabilizing effect of 9G when position 8 is mutated (i.e., not A). (H) Relative affinities for bound residues 1–9 (corresponding to PUM2 repeats 1′–9′) determined by global fit to the final model (additive nonconsecutive with coupling; Table 1A, Figure 3G). Error bars indicate 95% confidence intervals of the median, determined by bootstrapping analysis (Methods). (I) Relative affinities of A-specific PUM2 repeats. (J) Relative affinities of PUM2 repeats with conserved U-specific residues.

**Figure S5.**
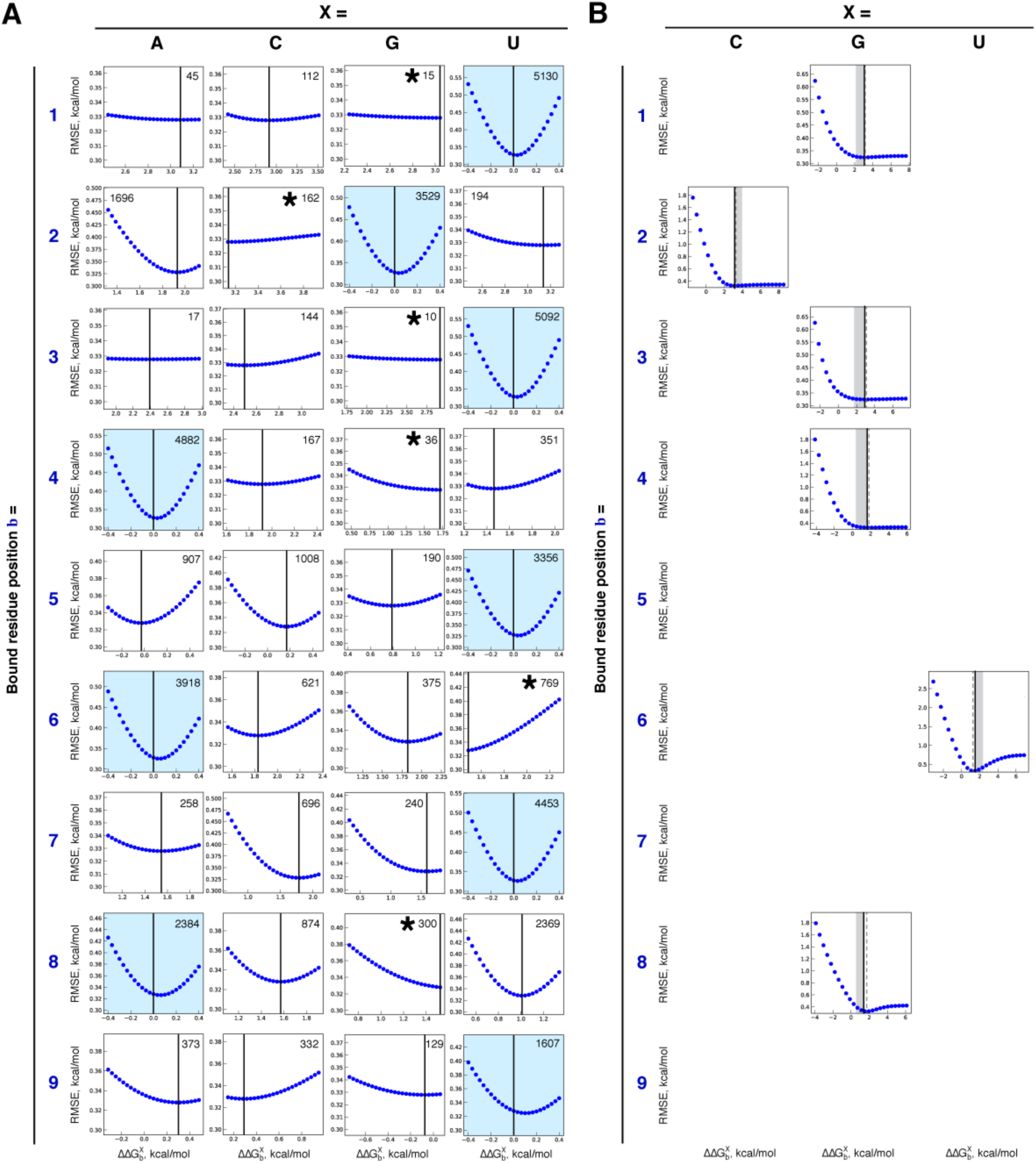
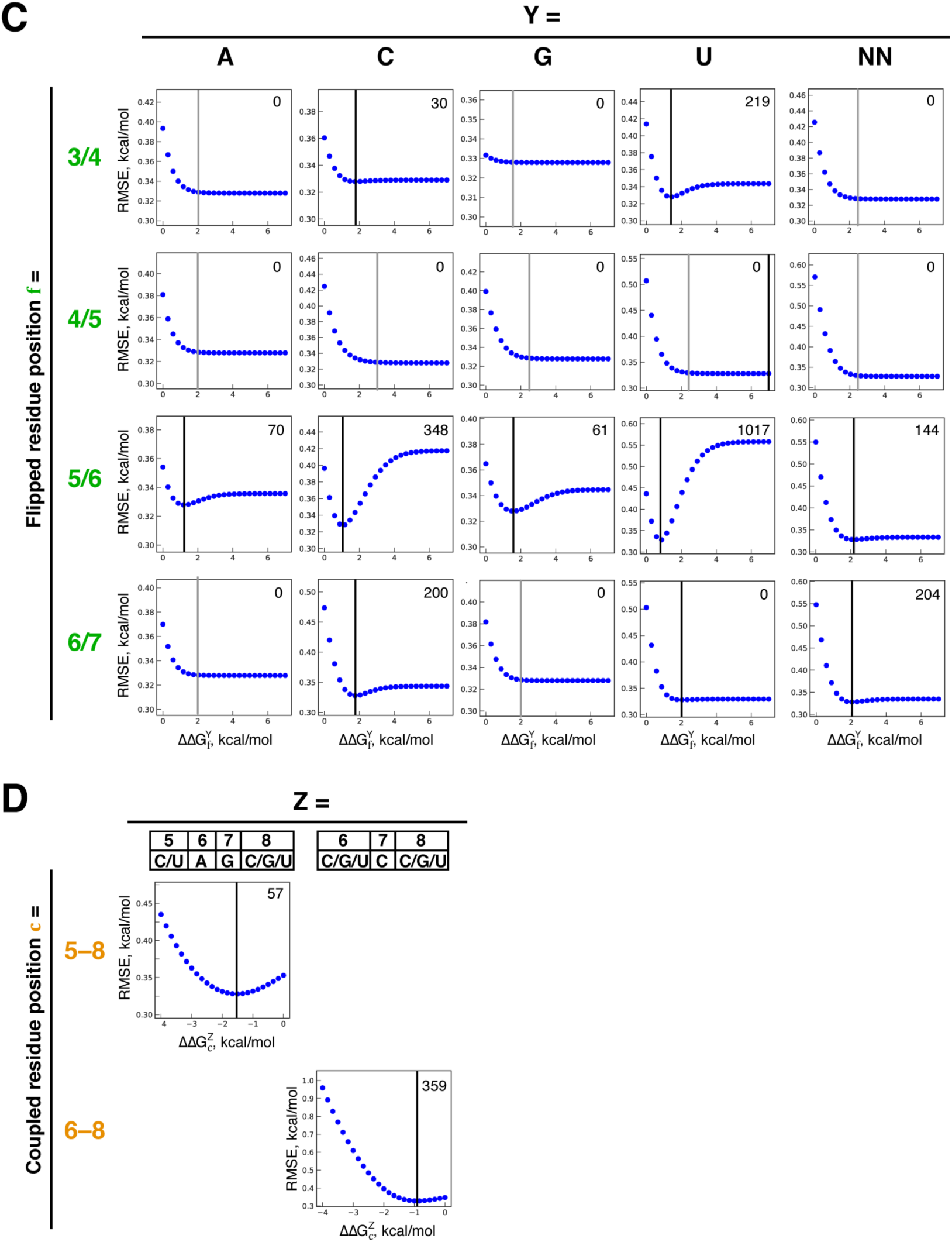
RMSE Dependence on Individual Fit Parameters in the Additive Nonconsecutive Model Including Coupling Terms, Related to Figure 3. (A) Bound residue terms (‘b’ in Figure 3G, Table 1A). The sensitivity of the global fit to each model parameter was individually assessed by varying the ΔΔG value across the indicated range, with the values of other parameters kept constant. ΔΔG values were constrained to measured single mutant penalties (median ± 0.4 kcal/mol) or to the 95% confidence intervals of the measured median if they exceeded 0.4 kcal/mol (Figure 2E and Table S2). Vertical black lines indicate the fit values. Asterisks indicate parameters for which the fit value was at the limit of the allowed range (see part B). The number of constructs containing each parameter in the best register (and any additional registers within 0.4 kcal/mol from the best one) is indicated. The slight offset from 0 observed for the consensus residues appears to be due to residual RNA structure (Methods). (B) Six binding terms for which the best fit value was at the limit of the allowed range (asterisks in part A) were refit without constraints on their ΔΔG value. The gray area indicates the original constraints. The solid lines indicate the original fit value, and the dashed line indicates the fit value after removing the constraints. In all cases, removing the constraints resulted in fit values within 0.2 kcal/mol of the original fit value. (C) Flipping terms (‘f’ in Figure 3G, Table 1B). As in part *A*, the sensitivity of the global fit RMSE to each term was assessed by varying the ΔΔG value across the indicated range, while the values of other parameters were kept constant. Black lines indicate the fit values; gray lines indicate lower limits for terms whose upper limits were undefined because these parameters were never featured in the most stable binding register (largely as a result of energetic redundancy between flipped positions). ‘NN’ indicates 2-nt flips of any sequence. (D) Coupling terms. RMSE sensitivity was assessed as in parts A–C. The 8/9 coupling term is not separately shown, as its value is determined by the position 9 binding terms (*A*; Table 1).

**Figure S6.**
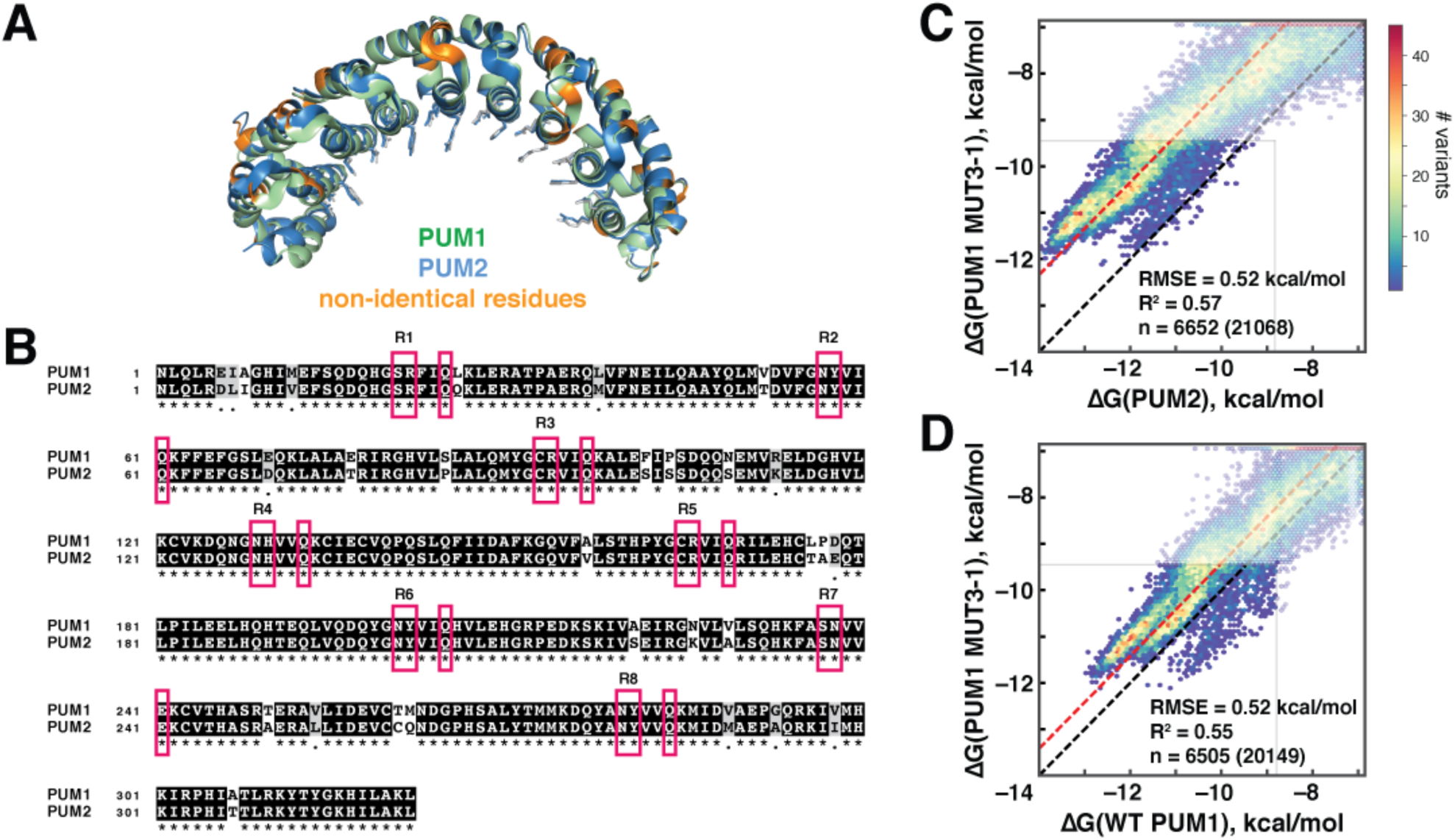
Comparison of PUM2 and Wild-Type and Engineered PUM1 Proteins, Related to Figure 5. (A) Structure alignment of the conserved RNA-binding domains of PUM1 (green; PDB ID: 3Q0L) and PUM2 (blue; 3Q0Q)(Lu and Hall, 2011). Residues that differ between the two proteins (n = 29/322; 9.0% of RBD residues) are highlighted in orange and are all positioned away from the RNA-contacting residues, which are shown in stick representation. The structural alignment was performed in PyMol (The PyMOL Molecular Graphics System, Version 1.8.6.2 Schrödinger, LLC.). (B) Sequence alignment of the RNA-binding domains of PUM1 and PUM2. Residues that form sequence-specific RNA interactions in each repeat are boxed in red. The sequence alignment was performed using T-Coffee and visualized using BOXSHADE (Notredame et al., 2000). (C) Scatterplot of the affinities of wild type and mutant PUM1. Black line indicates identical binding and red line indicates mean offset. (D) Comparison of mutant PUM1 and wild type PUM2, lines same as in part C.

**Figure S7.**
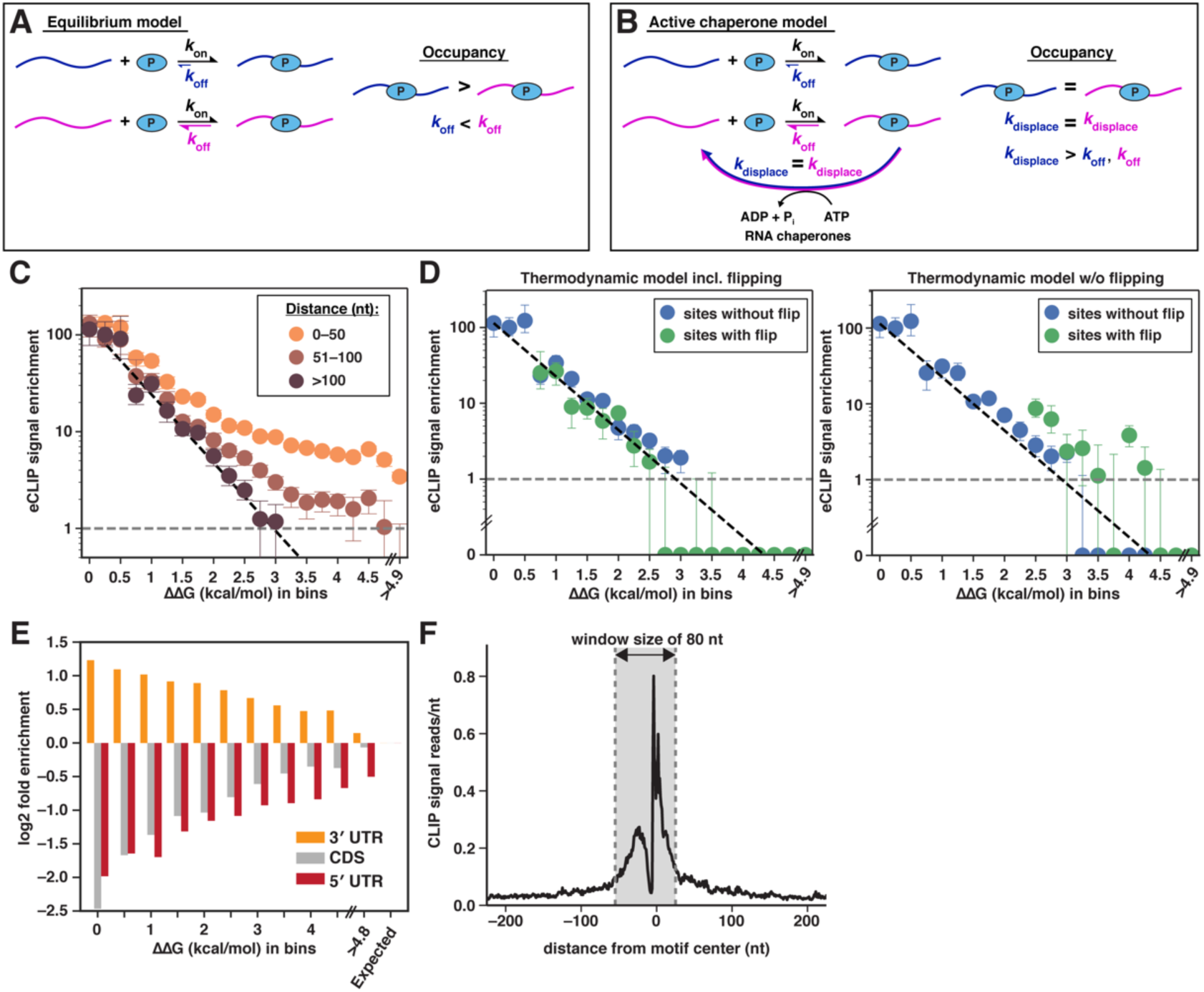
Assessment of PUM2/RNA Interactions in Vivo, Related to Figure 6. (A,B) Schematic representation of models for RNA occupancy in vivo in which active RBP (‘P’) displacement in vivo (e.g., by ATP-dependent RNA chaperones) yields identical occupancies for targets with different affinities. (A) Equilibrium model: Two targets with different affinities are shown, in blue and purple. The association rate constants (*k*_on_) for RBP binding to the two RNAs are identical, and the affinity difference is reflected in the greater dissociation rate constant (*k*_off_) for the second target (as is common and is observed for PUM2; data not shown). The occupancy (right) is greater for the first target due to its lower *K*_D_ value (*K*_D_ = *k*_off_/*k*_on_). (B) Kinetic model: If the rate constant of active displacement (*k*_displace_) is greater than either of the intrinsic dissociation rate constants, the occupancies are equal and independent of the affinity for the RNA targets (assuming equal *k*_on_ values). (C) Scatterplot comparing the median eCLIP enrichment across sites in bins of predicted relative affinity. Colors indicate sites with a nearby UGUA sequence within 50 nt, between 50–100 nt, or greater than 100 nt from the predicted site. The black dashed line indicates the expected change in the eCLIP signal for different predicted ΔΔG values, relative to the eCLIP signal in the lowest ΔΔG bin. Grey dotted line indicates expected background enrichment (= 1). Error bars indicate the 95% confidence intervals on the median. (D) Scatterplots comparing the median eCLIP enrichment across sites in bins of predicted ΔΔG, using either the full thermodynamic model (left) or a model that does not take into account flipped residues (right). Colors indicate sites predicted to have flipped residues (green) or not (blue). Only bins with at least 25 sites are shown to reduce noise. (E) Bars indicate the fold difference (log2) of the observed fraction of sites with the given annotation (5´ UTR, CDS, 3´ UTR) versus the expected fraction (based on the fraction of sites with ΔΔG > 4.5 kcal/mol). (F) Average number of eCLIP read stops per nucleotide (nt) around 3,703 PUM2 consensus binding sites (ΔΔG_pred_ < 0.5 kcal/mol) within annotated 3´ UTR, CDS, or 5´ UTR of expressed genes in K562 cells. The grey area indicates the interval relative to the binding site over which any eCLIP read stop was included in the calculation of eCLIP enrichment (80 nt window, offset by 15 nt relative to the center of the binding site).

## Methods

### Library design

Summary of library designs and complete sequence information is provided in Supplemental file 1.

### Library preparation

#### Ordering

DNA constructs consisting of the PUF library and short constant regions for subsequent PCR assembly (Figure S1D & Table S5; 5´–TGTATGGAAGACGTTCCTGGATCC–[Variable region]–AGATCGGAAGAGCGGTTCAG–3´) were ordered from CustomArray, Inc. as part of a 90,000 oligo pool of 130 nt sequences. Each of the 34,927 unique sequences in the library (including variants not discussed herein) was included at least in duplicate to increase the probability of error-free generation. In cases where the designed sequence was shorter than 130 nt, the construct was “padded” at the 3*′* end with a random sequence that was eliminated during PCR assembly. Primers and DNA oligonucleotides used in the RNA-MaP protocol were ordered from Integrated DNA Technologies (IDT).

#### Emulsion PCR

The oligonucleotide pool was amplified using emulsion PCR (ePCR) (Williams et al., 2006), allowing us to decrease length and other biases during PCR amplification of our highly diverse library (lengths of 64–130 nt, variable structure content). We closely followed a MYcroarray adaptation of the ePCR protocol from (Williams et al., 2006), as detailed below. Flat-bottom glass vials (1 mL) were cleaned with sterile water, dried, covered with parafilm, and frozen in a petri dish filled with sterile water. The oil phase was prepared from 4% (v/v) ABIL EM-90, 0.05% (v/v) Triton X-100 and 96% (v/v) mineral oil. The 50 μL aqueous phase consisted of 1.45 ng/μL of the CustomArray oligo pool, 0.2 mM of dNTP, 1x Phire II buffer, 1 μL of Phire Hot Start II DNA Polymerase (Thermo Fisher Scientific), 0.5 mg/mL BSA, and 2 μM of each of *RNAPstall* and *Read2* primers (Table S6 and Figure S1D). A 300 μL aliquot of the vortexed, pre-chilled oil phase was added to the glass vial embedded in the ice-filled petri dish and stirred on a stir plate with a sterile magnetic bar at 1000 rpm for 5 min. The aqueous phase was then added in five 10 μL aliquots and stirred for another 10 min. The emulsion was divided between seven PCR tubes and amplified using the following settings:

98 °C for 30 s

40 cycles of:

98 °C for 10 s

65 °C for 10 s

72 °C for 30 s

72 °C for 5 min.

Completed PCR reactions were pooled in a 1.7 mL Eppendorf tube, and 1 μL of gel loading dye was added to visualize the aqueous phase. Mineral oil (100 μL) was added and the mix was vortexed for 30 s, followed by centrifugation for 10 min at 13,000g. The oil was discarded and 1 mL diethyl ether was added, the mixture was vortexed in fume hood for 3 min and centrifuged for 1 min at 13,000 g. Diethyl ether was discarded, 1 mL ethyl acetate was added, and the mixture was vortexed in fume hood for 3 min and centrifuged for 1 min at 13,000 g. Ethyl acetate was removed and the diethyl ether extraction step was repeated, followed by discarding the diethyl ether. The tube was incubated for 5 min at 37 °C with an open cap to allow residual diethyl ether to evaporate. Water (40 μL) and Agencourt AMPure XP beads (Beckman Coulter; 72 μL) were added and incubated for 15 min at room temperature; the supernatant was removed, the beads were washed with 70% ethanol (2 x 100 μL), dried, and the DNA was eluted in 10.5 μL water.

#### Size fractionation

To further prevent bias toward short oligonucleotides during the subsequent PCR assembly steps, the ePCR-amplified library was fractionated by length on an 8% polyacrylamide gel. Following SYBR Green staining (1x; Lonza), the library-containing lane was covered with aluminum foil to prevent UV-induced damage (Grundemann and Schomig, 1996; Sinha and Hader, 2002), and divided into 6 fractions based on UV visualization of marker lanes. The cut-out bands were frozen on dry ice and eluted overnight in TE buffer (10 mM Tris-HCl, pH 8.0, 1 mM Na_2_EDTA) on a rotating platform at 8 °C. The DNA was purified using the Qiagen Gel Extraction Kit (using a protocol adapted for PAGE purification: http://www.qiagen.com/kr/resources/resourcedetail?id=1426dbb4-da09-487c-ae01-”c587c2be14c3&lang=en, with Qiagen MinElute columns). To remove residual co-purified short fragments, each fraction was re-purified on an 8% denaturing gel (8 M urea). For denaturing PAGE, the samples and a Low-MW DNA ladder (New England Biolabs; NEB) were heated in loading buffer (84% (v/v) formamide, 50 mM Na_2_EDTA, 0.04% Xylene cyanol, 0.04% Bromophenol blue; 2.8 μL loading buffer per 5 μL sample) at 90 °C for 3 min immediately before loading. The gel was stained with SYBR Gold, the library-containing lanes were covered with aluminum foil and fractions were cut out based on UV visualization of marker lanes. (Additional lanes with 83 nt and 129 nt DNA oligonucleotides were used to facilitate alignment of the NEB low-MW marker with desired lengths.) The DNA was extracted from the gel as above. Purified fractions were re-amplified using the *Read 2* and *RNAPstall_adapt* primers (primer sequences and PCR settings are indicated in Tables S6 & S7) and were purified using Qiagen MinElute PCR Purification Kit. In all cases here and below, the number of PCR cycles was determined by quantitative PCR (qPCR), using the same primer and template concentrations as in preparative PCR, but in the presence of 0.2–0.5x SYBR Green. To prevent accumulation of by-products, cycle numbers corresponding to about one-third saturation (C_t_ value) were used in preparative PCR reactions. Each library fraction was amplified for two to three different numbers of cycles around the C_t_ value, and only reactions lacking high-molecular weight byproducts were propagated to the next step.

#### PCR assembly

Each purified length fraction was assembled into the final RNA array construct with the *C_read1_bc_RNAP, D_read2, C_adapter* and *D_adapter* primers, as illustrated in Figure S1D (see Tables S6 & S7 for primers and conditions). The *C_read1_bc_RNAP* primer contained a randomized 15 nt ‘barcode’ region that served as a unique molecular identifier (UMI) and allowed high-confidence sequence mapping during subsequent steps (Buenrostro et al., 2014). The PCR products were purified using QIAquick PCR purification kit (Qiagen).

#### Bottlenecking

To ensure that multiple copies of each UMI were present on the RNA array, the library was bottlenecked to ∼700,000 total molecules (Buenrostro et al., 2014; Denny et al., 2018; Kivioja et al., 2012). UMI redundancy allows distinguishing between sequencing errors and real sequence differences, as errors are unlikely to co-occur in both the UMI and the variable region (see **Computational analyses** below). To bottleneck, the PCR products were quantified by qPCR relative to the PhiX standard (Illumina). As noted above, the six ‘sublibraries’, corresponding to the different oligonucleotide lengths in our library, were kept separate during all pre-sequencing PCR steps to minimize bias in the final library assayed by RNA-MaP. Dilutions of 1000-fold and 10,000-fold for each fraction were prepared in 0.1% Tween 20. The PhiX standard (Illumina) was diluted to 200 pM and seven serial dilutions were prepared in 0.1% Tween 20. The DNA was then added to a PCR master mix containing 500 nM *OligoC* and *OligoD* primers, 200 μM dNTP mix, 0.5x SYBR Green, 3% DMSO, 0.02 U/μL Phusion DNA Polymerase (Thermo Fisher Scientific), and 1x Phusion buffer. The cycling conditions were: 98 °C for 30 s, followed by 35 cycles of 98°C for 10 s, 63 °C for 30 s, and 72 °C for 30 s. The library concentrations were determined based on the PhiX standard curve of C_T_ values over concentration (determined in duplicate). The volumes corresponding to a total of 700,000 molecules across all sublibraries were calculated, and each sublibrary was amplified with *OligoC* and *OligoD* primers. The PCR reactions were purified with QIAquick PCR cleanup kit (Qiagen), concentrations of 1000-fold dilutions were quantified by qPCR and the different fractions were combined for sequencing. Due to a short OligoD byproduct detected as dominant species in the initial sequencing of our library, the bottlenecked library fractions were re-purified on a denaturing 8% acrylamide gel and amplified using Phusion Hot Start II DNA Polymerase (Thermo Fisher Scientific) instead of regular Phusion, which eliminated the primer byproduct. This library was sequenced and used in all RNA-MaP experiments reported herein.

### RNA array preparation

The bottlenecked, qPCR–quantified library fractions were combined and sequenced using MiSeq Reagent Kit v3 (150-cycle). To ensure appropriate density of RNA clusters in the RNA-MaP experiments, our library constituted 9–15% of the total 6–9.6 fmol DNA. The remaining DNA consisted of 84–90% PhiX DNA and 1% of a fiduciary marker oligonucleotide (Table S8) that was used for image alignment in binding experiments. The final numbers of transcribable clusters were 3.6×10^5^ and 6.5×10^5^ on the sequencing chips used in this study.

### Protein expression and purification

The RNA-binding domains of *H. sapiens* PUM1 (828-1176; isoform 2), PUM2 (706-1059; isoform 1), and mutant PUM1 (MUT3-1 in (Cheong and Hall, 2006)) (828-1176) were cloned into a custom pET28a-based expression vector in frame with an N-terminal His-tag and a SNAP tag (New England Biolabs) at either the N- (PUM2) or C-terminus (hPUM1 and hPUM1 MUT3-1; primers and plasmid sequences available upon request). Constructs were sequenced and transformed into *E. coli* protein expression strains BL21 (DE3) or RIPL BL21 CodonPlus (Agilent) cells. Protein expression was induced at an OD600 of between 0.6–0.8 with 0.5–1 mM IPTG at 18–20 °C for 18–20 h. Cell pellets were lysed four times using an Emulsiflex (Avestin) in Buffer A containing 20 mM Na-HEPES, pH 7.4, 500 mM potassium acetate (KOAc), 5% glycerol, 0.2% Tween-20, 10 mM imidazole, 2 mM DTT, 1 mM phenylmethylsulfonyl fluoride (PMSF) and 2X Complete Mini protease inhibitor cocktail (Roche). The lysate was centrifuged at 20,000 g for 20 min to remove membranes and unlysed cells. Nucleic acids in the lysate were precipitated with dropwise addition of Polyethylene Imine (Sigma) to a final concentration of 0.21% (v:v) with constant stirring at 4 °C and pelleted by centrifugation at 20,000g for 20 min. Cleared lysates were then loaded on a Nickel-chelating HisTrap HP column (GE), washed extensively, and His-tagged proteins were eluted over a 10–500 mM imidazole gradient. Protein fractions were pooled and desalted into Buffer B (20 mM Na-HEPES, pH 7.4, 50 mM KOAc, 5% glycerol, 0.1% Tween-20, 2 mM DTT) using a HiPrep 26/10 desalting column. The His-tag was removed by incubation with TEV protease for 13–16 h at 4 °C, and the protein solution was loaded for a second time on the HisTrap HP column. The flow-through containing cleaved protein was collected and subsequently desalted into Buffer B. The protein was then loaded on a Heparin or HiQ column and eluted over a linear gradient of KOAc from 50 to 1000 mM. Fractions were pooled and desalted into Buffer C containing 20 mM Na-HEPES, pH 7.4, 100 mM KOAc, 5% glycerol, 0.1% Tween-20 and 2 mM DTT, concentrated using Millipore 10K filters and diluted two-fold with Buffer C containing 80% glycerol for final storage at –20 °C.

### Cy3B-labeling of SNAP-tagged proteins

Cy3B-labeled SNAP tag substrate was prepared by coupling Cy3B NHS ester (GE Healthcare,0.75 μmol) with 1.5-fold excess (1.13 μmol) of amine-terminated benzylguanine (NH2-BG; New England BioLabs) in the presence of 1.13 μmol triethylamine in dimethylformamide. The reaction (103 μL) was incubated overnight on a rotating platform at 30 °C. The Cy3B-BG product was purified by reverse phase HPLC on an Agilent ZORBAX Eclipse Plus 95Å column and dried by speed-vac evaporation (46% yield).

SNAP-tagged PUF proteins were labeled by incubating 5–10 μM of purified protein with 20 μM of Cy3B-BG in Buffer C. The tube was covered with aluminum foil and rotated at 4 °C for 12–14 h. Unincorporated dye was removed with Zeba Spin Desalting Columns (Thermo Fisher Scientific) equilibrated with Buffer C; the protein was concentrated using Millipore 10K filters and diluted two-fold with Buffer C containing 80% glycerol for final storage at –20 °C. The labeling efficiencies (based on total protein concentration and Cy3B absorbance at 559 nm; Cy3B extinction coefficient: 130,000 M^−1^cm^−1^) were 60% (PUM2-SNAP), 53% (SNAP-PUM1) and 36% (mutant SNAP-PUM1).

### RNA-MaP measurements

#### Imaging station setup

The RNA-MaP imaging platform was built out of a repurposed Illumina GAIIx instrument with custom-designed additions as described in (Buenrostro et al., 2014) (Denny et al., 2018; She et al., 2017). Briefly, the custom additions included a fluidics adapter interface to pump reagents to the MiSeq flow cell, a Peltier-based temperature-controlled platform to house the flow cell, an autosampler with 96-well cooling block for RNA-MaP reagents, and a dual–color laser excitation system. Two lasers were employed: a 660 nm ‘red’ laser with a 664 nm long pass filter and a 530 nm ‘green’ laser with a 590 nm band pass filter. Matlab scripts developed in-house were used to control the fluidics, temperature, position, and imaging of the flow cell. Flow cell images were acquired with 400 ms exposures at 200 mW laser power. Camera focal distances were determined through iterative rounds of imaging of the flow cell and adjustment of the camera’s z-position.

#### RNA transcription in the flow cell

Using the imaging station fluidics system, the flow cell was washed with 5 mM Na_2_EDTA in formamide to remove hybridized DNA (250 μL flowed at 100μL/min, 55 °C), followed by Reducing buffer (100 mM Tris•HCl, 125 mM NaCl, 0.05% Tween 20, 100 mM Tris[2-Carboxyethyl]phosphine-HCl (TCEP), pH 7.4) to remove any residual fluorescence from the sequencing reaction (390 μL, 10 min at 60 °C). A fluorescent probe complementary to the RNA Polymerase stall sequence (*Fluorescent_stall*´; sequences of oligonucleotides used in the RNA-MaP protocol are indicated in Table S8) was then annealed to the library and imaged to determine the efficiency of the cleaning steps (500 nM *Fluorescent_stall*´ in Annealing buffer: 1x SSC buffer, 7 mM MgCl_2_, 0.01% Tween 20; 11 min at 37 °C). After imaging, the fluorescent probe was removed by washing with 250 μL of 100% formamide (55 °C). The flow cell was washed with Wash buffer between steps (290 μL; 10 mM Tris•HCl, pH 8.0, 5 mM Na_2_EDTA, pH 8.0, 0.05% Tween 20). Henceforth, wash steps were performed with a 250 μL volume of the specified buffer, unless otherwise noted.

To prepare double-stranded DNA (dsDNA), 5′–biotinylated primer (*Biotin_D_Read2*, 500 nM) was annealed to the library in Hybridization buffer (5x SSC buffer, 5 mM Na_2_EDTA, 0.05% Tween 20) for 15 min at 60 °C followed by a 10 min incubation at 40 °C. The fluorescent oligonucleotide complementary to the fiducial marker (*Fiducial_flow*) was also included in the hybridization mixture at 250 nM. After washing the flow cell with Annealing buffer, an additional 500 nM of *Biotin_D_Read2* (and 250 nM *Fiducial_flow*) was annealed to the library in Annealing buffer at 37°C for 8 min. The flow cell was then washed with Klenow buffer (1x NEB buffer 2 (NEB B7002S), 250 μM each dNTP, 0.01% Tween 20). Double-stranded DNA was generated by pumping 130 μL of 0.1 U/μl Klenow fragment (3′-5′ exo(–); NEB M0212L) into the flow cell in three stages separated by 10 min intervals each. The flow cell was maintained at 37 °C for this period. Unextended single-stranded DNA templates were subsequently blocked by annealing a non-fluorescent version of the stall probe (*Dark_stall*′) in a process identical to the one described above.

After dsDNA generation, 1 μM streptavidin in Annealing buffer was pumped into the flow cell and allowed to bind to the biotinylated primer for 5 min at 37 °C. Excess streptavidin was then washed out of the flow cell with Annealing buffer. Unbound biotin binding sites in the streptavidin tetramer were saturated by incubating the flow cell for 5 min with 5 μM free biotin in Annealing buffer. Excess unbound biotin was washed out with Annealing buffer.

RNA transcription proceeded in two stages, initiation/stall and extension. In the initiation/stall phase, 130 μL of 0.06 U/μL *E. coli* RNA polymerase holoenzyme (RNAP; NEB M0551S) was allowed to initiate transcription for 20 min at 37 °C on the dsDNA templates in Initiation buffer, which lacked CTP (20 mM Tris•HCl pH 8.0, 7 mM MgCl_2_, 20 mM NaCl, 0.1% BME, 0.1 mM Na_2_EDTA, 1.5% glycerol, 0.02 mg/mL BSA, 0.01% Tween 20, and 2.5 μM each of ATP, GTP, and UTP). Upon encountering the first cytosine (C27), the polymerase stalls, thereby sterically preventing the loading of additional enzymes on the same template (Buenrostro et al., 2014). Excess RNAP was washed out of the flow cell with Initiation buffer. Subsequently, Extension buffer was added, which contained all 4 ribonucleotides (20 mM Tris•HCl pH 8.0, 7 mM MgCl_2_, 20 mM NaCl, 0.1% BME, 0.1 mM EDTA, 1.5% glycerol, 0.02 mg/mL BSA, 0.01% Tween 20, and 1 mM each of ATP, GTP, UTP and CTP). The Extension buffer also contained 500 nM each of *Fluorescent_stall*′ and *Dark_read2* oligonucleotides, which were intended to block the regions flanking the variable region in the nascent RNA transcript (Fig. 1C) and prevent undesired intramolecular interactions and also allow visualization of the transcript. Transcription was allowed to proceed for 10 min at 37 °C. RNA polymerase eventually is stalled at the streptavidin roadblock at the end of the DNA template, exposing the nascent RNA molecules for binding experiments (Fig. 1C).

To ensure complete blocking of RNA regions flanking the variable sequence, transcription was followed by further hybridization of *Fluorescent_stall*′ and *Dark_read2* oligonucleotides (500 nM) for 10 min at 37 °C in Annealing buffer. Finally, the flow cell was washed with Binding buffer (20 mM Na-HEPES, pH 7.4, 100 mM KOAc, 0.1% Tween-20, 5% glycerol, 0.1 mg/ml BSA, 2 mM MgCl_2_ and 2 mM DTT), the temperature was lowered to 25 °C (except for 37 °C experiments), and the flow cell was imaged to quantify the fluorescence from the RNA-annealed *Fluorescent_stall*′ probe.

#### RNA-MaP equilibrium binding experiments

To determine PUM1/2 binding affinities, the RNA library was sequentially equilibrated with increasing concentrations of Cy3B-labeled PUM proteins, and the amount of Cy3B fluorescence colocalized with each RNA cluster was determined at each concentration. Two-fold serial protein dilutions (15–17) were prepared in 1x Binding buffer and were stored in light-protected tubes on ice or in the 4 °C autosampler chilling block until the incubation. Protein solution (460 μL) was pumped into the flow cell at each concentration and incubated for times ranging from 33 min for the lowest concentrations to 19 min for the highest protein concentrations (25 °C; 15–23 min at 37 °C). These incubation times were established to be sufficient for equilibration by association and dissociation time courses (halftime ≤ 5.3 min; see also (Vaidyanathan et al., 2017)). The incubation temperature was 25 or 37 °C, as indicated for the individual experiments.

### Computational analyses

#### Processing sequencing data

Illumina MiSeq sequencing data were computationally processed to extract the tile identifier and the x- and y-locations of each sequenced cluster from the fastq file output (Denny et al., 2018; She et al., 2017). Sequence clusters were divided into three categories: (1) clusters encoding our RNA library, (2) clusters containing the fiducial sequence, and (3) inert “background” sequence clusters lacking the RNAP initiation site or the fiducial sequence. This assessment was based on alignment of the read1 sequence to (1) the RNA polymerase initiation site and stall sequence “TTTATGCTATAATTATTTCATGTAGTAAGGAGGTTGTATGGAAGACGTTCCTGGATCC”, or to (2) the fiducial sequence “CTTGGGTCCACAGGACACTCGTTGCTTTCC”, or (3) neither, respectively.

While every cluster was fit during the 2D Gaussian fitting (described below), only clusters containing the fiducial sequence were used during the cross-correlation of the images to the sequencing data. *K*_D_ fitting was only performed on RNA-encoding clusters (described below).

#### Fitting images

To attribute fluorescent binding events to individual sequence variants, the images taken during the RNA array experiment were mapped to the sequencing data output from the Illumina MiSeq. Each image had a set of fiducial clusters, which were visualized with a fluorescently labeled complementary oligo (see *RNA transcription in the flow cell* above). The x- and y-locations of each fiducial cluster were cross-correlated with each fluorescent image in an iterative fashion to determine a smooth function of x and y that represents the location-dependent offset between the image and the sequencing data locations. This “registration offset map” enabled correlation between image and sequencing locations at sub-pixel resolution, as described in (She et al., 2017). Once this map was determined, each image was fit to the sum of 2D Gaussians, with each Gaussian centered at each of the cluster locations from the registered sequencing data output. The quantified fluorescence of each cluster was thus the integrated fluorescence within the fit 2D Gaussian (*f* = 2*πaσ*^2^ where *f* is the integrated fluorescence and *a* and *σ* are the fit amplitude and standard deviation, respectively).

#### Identifying library variants from sequencing data

Incorporation of a 15-nt unique molecular identifier (UMI) in our library allowed us to minimize the effect of sequencing errors when associating each sequenced cluster with its underlying sequence variant, as described in (Buenrostro et al., 2014). Prior to sequencing, the library was bottlenecked and reamplified (see **Library preparation** above), resulting in increased representation of each UMI that survived the bottlenecking. We assumed that all sequences associated with the same UMI came from the same molecular variant, so that any variation between these sequences was the result of sequencing error. To resolve sequencing errors, a consensus sequence of the associated library variant was determined for each UMI with a per-base voting strategy. Only UMIs with a significant fraction of variants matching the consensus sequence were used, as evaluated by a binomial null model. For each UMI, a p-value was calculated based on the number of associated sequences that matched the consensus sequence and the total number of sequences, assuming a rate of success under the null model of 25%. UMIs with a higher rate of matching than expected by chance under the null model (i.e., with p-value < 0.01) were defined as successfully associating with a consensus sequence. Clusters were then associated to a designed library variant based on the cluster’s UMI.

#### Fluorescence normalization

To account for inter-cluster variation in maximum fluorescence, we normalized the amount of protein bound at a given cluster by the total amount of transcribed RNA in that cluster. This normalization was performed by dividing the integrated fluorescence of the cluster in the green channel (i.e., the channel imaging the bound protein) by the integrated fluorescence of the same cluster in the red channel (i.e., the channel imaging the fluorescent oligo annealed to the transcribed RNA). To prevent dividing by small numbers and inflating the normalized signal towards infinity, values of the red channel fluorescence below the threshold of the first percentile of the distribution of the red channel fluorescence across clusters were set to the value of the threshold.

#### Determining the free energy of binding

The normalized fluorescence values of bound protein across different solution protein concentrations were used to determine the equilibrium dissociation constant (*K*_D_) between the protein and each RNA variant. The fitting procedure was split into several steps to allow robust fitting across a range of affinities, and the binding model accounted for observed non-specific binding events. In brief, the normalized fluorescence values for each individual cluster were fit to a binding curve to obtain best-fit values for *K*_*D*_ and other fit parameters (see *Single cluster fitting* below). These best-fit values for individual clusters were used to determine distributions for fit parameters that we expect to be variant-independent —i.e., *K*_D,NS_, *f*_*min*_, and *f*_*max*_, which are each defined below. The distributions of these common values across library variants were used to refine the *K*_D_ value for each RNA variant.

#### Single cluster fitting

Initially, fluorescent values for each cluster were fit to a binding curve. This binding model incorporated a nonspecific binding term, as follows:

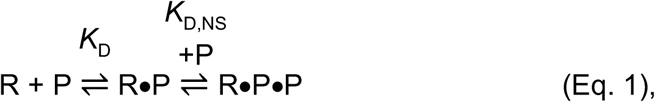

where R is RNA, P is protein, *K*_D_ is the dissociation constant (*K*_D_ = e^ΔG/RT^) and *K*_D,NS_ is the non-specific dissociation constant for a second protein monomer that binds to the RNA/protein complex 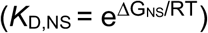.

The normalized fluorescence of a cluster at protein concentration [P] can be defined as:

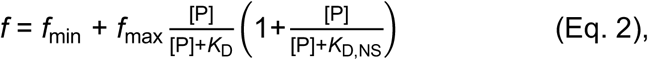

where *f*_min_ is the background fluorescence in the absence of bound protein, *f*_max_ is the fluorescence signal at saturation, and [P] is the concentration of the protein in solution (here, [P] ≈ [P]_total_). This model was used to account for an observed increase in fluorescence at high concentrations of protein after apparent saturation (Figure 1D). We did not observe a corresponding increase in fluorescence for non-binding variants, and the extent of ‘non-specific’ binding increased with greater specific binding affinity. Together, these observations support the model that a second protein monomer binds to the RNA-protein complex on chip (as opposed to non-specific binding to the DNA or unbound RNA, which would be independent of bound protein).

Least squares fitting was performed using the Python package lmfit (v 0.8.3). The initial estimates and constraints are as follows: *f*_*min*_ was initialized to the median fluorescence across clusters in the images with no protein applied, and was constrained to be not less than zero during the fitting; *f*_*max*_ was initialized at the maximum fluorescence observed at any concentration of protein for that cluster, and was similarly constrained to not be less than zero; *K*_D_ was initialized at the highest concentration of the protein; and *K*_D,NS_ value was initialized at five-fold the highest concentration of the protein.

#### *Finding common values for* K*_D,NS_,* f*_min_, and* f*_max_ across variants*

Allowing all four free parameters (*f*_min_, *f*_max_, *K*_D_, and *K*_D,NS_) to float during the fitting process led to some spurious effects. In particular, variants with low affinity that do not achieve saturation within the probed protein concentration range can be fit approximately equally well with different values for *f*_max_ and *K*_D,NS_, ultimately leading to uncertainty in the fit value for *K*_D_. For example, a variant that does not achieve saturation may be fit equally well with lower values for *K*_D_ and *f*_max_, higher values for *K*_D_ and *f*_max_, or a higher value for the *K*_D,NS_ and lower value for *K*_D_. On the other hand, the library contains numerous tightly bound variants which have achieved saturation and from which we can extract most likely values for the sequence-independent parameters *f*_min_, *f*_max_, and *K*_D,NS_. These values are largely constant across different molecular variants that did achieve near-saturation (Figure S1E,F), allowing us to reasonably assume that the same values are applicable to all molecular variants, i.e., even those that did not achieve saturation, and applying these well-defined estimates for *f*_min_, *f*_max_, and *K*_D,NS_ allows more confident fitting of the *K*_D_ values. To limit noise, the estimates for *f*_min_, *f*_max_, and *K*_D,NS_ were determined based on the per-variant values of each fit parameter, where per-variant values are the median of the single-cluster values associated with the same molecular variant.

#### *Estimating* f*_min_*

The value for *f*_min_ was largely consistent across variants (Figure S1E); thus, the estimate for this fit parameter was simply the median value across variants.

#### Estimating the distribution of f_max_

To define the distribution of *f*_max_ values across molecular variants, a subset of variants with low and precisely measured *K*_D_ values was used, based on the single cluster fits. Variants used to define this distribution had *K*_D_ values less than 5% of the highest concentration of protein. In addition, the precision of the per-variant values of *K*_D_ was evaluated based on the proportion of the variant’s single cluster fits having a goodness-of-fit (R^2^) greater than 0.5, the standard error on ΔG (ΔG = RTln(*K*_D_)) less than 1 kcal/mol, and the standard error on the fit *f*_max_ less than *f*_max_. If a significant fraction of the clusters associated with this variant passed these filters, then the variant was considered to have a “precise” measurement of *K*_D_. Significance was assessed based on rejecting the null hypothesis that 25% of clusters would pass all these filters by chance alone (binomial p-value < 0.01. For variants that did not achieve saturation the *f*_max_ was undefined, so the distribution of *f*_max_ values across the tightly bound molecular variants was used to find error estimates on the *K*_D_ values that reflect this uncertainty.

The *f*_max_ distribution across these variants was fit to a gamma distribution with fixed mean for the entire experiment, but whose standard deviation was dependent on the number of clusters per molecular variant (i.e., standard deviation should be proportional to 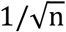, where n is the number of clusters per variant). This distribution reflects the fact that as the number of clusters increases, we can obtain more precise estimates of *f*_max_ and thus *K*_D_. The mean *f*_max_ value was obtained by fitting the per-variant *f*_max_ values to a gamma distribution, and obtaining the mean of the distribution, μ _global_. To obtain the standard deviation of the *f*_max_ distribution at each value of *n* (where *n* is the number of clusters per variant), the distribution of per-variant *f*_max_ values of variants with *n* clusters was fit to the gamma function *f*(*x*), for every normalized fluorescence value *x*:

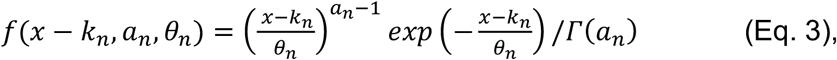

where *k_n_* is a free parameter, *a_n_ θ_n_* = *μ _global_*, and the resulting standard deviation is 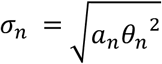. Allowing *k* to float resulted in better estimates of the standard deviation when distributions were more asymmetric, as often occurred for variants with small *n*. The value of −_n_ was initialized at 0, and σ_n_ was initialized at the standard deviation of the *f*_max_ values.

The values for σ_n_ may be subject to noise, given that some values of *n* had many more variants associated with that number of clusters than others did. To smooth these values, the σ_n_ values were used to fit the expected analytical function that defines the relationship between number of measurements and standard error, 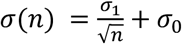, where σ_0_ and σ_1,_ are free parameters. σ_,1_ is the standard deviation on the estimate of *f*_*max*_ with only one measurement, and σ_0_ represents the standard deviation of *f*_max_ among different molecular variants if all variants were measured an infinite number of times. We expect this term to be nonzero in the case that the *f*_max_ depends on the molecular variant: e.g., if certain variants are trapped in stable secondary structures that do not unfold on the time scale of the binding experiment.

The estimator for *f*_max_ for each molecular variant with n clusters per variant is then the gamma distribution:

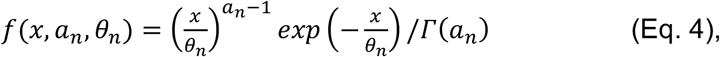

where 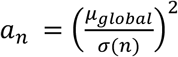 and 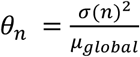 which depends only on *μ_global_* and σ(*n*), and the number of clusters per variant *n*.

#### *Estimating* K_D,NS_

The value for *K*_D,NS_ was determined by taking the median value across the subset of variants with low and precisely measured *K*_D_ values, based on the single cluster fits, as described above for determining the distribution of *f*_max_ values. The ΔG_NS_ terms (= RTlog(*K*_D,NS_)) for each protein were similar, with the following values: PUM2: –8.98 and –8.87 kcal/mol for replicates 1 and 2, respectively (25 °C), –8.65 kcal/mol (37 °C); WT PUM1: –8.46 kcal/mol; mutant PUM1: –8.19 kcal/mol.

#### *Applying common values for* K*_D,NS_,* f*_min_, and* f*_max_ to refine estimates for* K*_D_*

Binding isotherms were refined for all variants using the variant-independent values for *f*_min_, *K*_D,NS_, and, for the cases in which the variant did not achieve near-saturation, *f*_max_. To perform this refinement, clusters associated with each variant were resampled to obtain median fluorescence values across resampled clusters. This vector of median fluorescence values was fit to a binding isotherm with the values for *f*_min_ and *K*_D,NS_ fixed to the variant-independent values obtained above. For the cases where the variant did not reach near-saturation the value for *f*_max_ was also fixed. Not achieving near-saturation was defined as the median fluorescence value at the highest concentration of protein being less than the lower bound of the 95% confidence interval on *f*_max_. In this case, the value for *f*_max_ was sampled from the variant-independent distribution of *f*_max,n_ (with *n* equal to the number of clusters associated with this variant) (Eq. 4). To obtain uncertainty estimates on the fit *f*_max_ and *K*_D_ values, this resampling procedure was repeated 100 times. In the case that the variant did not achieve saturation, a different value for *f*_max_ was sampled for each iteration.

The 95% confidence intervals on *K*_D_ were obtained using these 100 values. The median fit *K*_D_ obtained from the initial single cluster fits was used as the initial value in the least squares fitting of each fitting iteration. For variants where *f*_max_ was allowed to float, the median fit *f*_max_ was used as the initial value.

#### Fitting background clusters to determine maximum measurable ΔG

To obtain a reasonable estimation of the highest *K*_D_ that can be measured by this method, we applied this fitting procedure to a set of “background” variants––i.e., variants on the chip lacking an RNAP initiation site. To normalize the bound fluorescence in the green channel to a similar scale as those clusters that do transcribe RNA, the fluorescence values were divided by the median fluorescence value in the red channel across clusters that do have RNAP initiation sites. Background clusters were randomly assigned to a “variant ID”, such that the set of “background variants” had a similar number of associated clusters as our library members. Finally, fitting was carried out as described in “*Applying common values for* K*_D,NS_,* f*_min_, and* f*_max_ to refine estimates for* K_*D*_”, with the variant-independent values for f_*min*_, f_*max*_, and K_D,NS_ applied. The reliable ΔG threshold determined by this analysis for PUM2 was approximately –8.5 kcal/mol (Figure S1F), and only variants with ΔG values less than this threshold (corrected for active protein fraction) were included in the high-confidence affinity data reported herein.

### Data filtering

Variants were included in our analyses if they met the following criteria (unless otherwise indicated): (1) observed ΔG values lower than –8.5 kcal/mol; this range was established as clearly distinguishable from background of non-transcribed clusters (see “Fitting background clusters to determine maximum measurable ΔG” above); (2) five or more clusters in at least one replicate experiment, to allow robust cluster statistics; in cases where one of the replicates contained fewer than five clusters, only the ΔG value from the replicate with 5 or more clusters was used; (3) 95% bootstrap confidence interval of the ΔG value (or weighted mean of replicate ΔG values) less than 1 kcal/mol; all 95% CI values were corrected to account for inter-experimental error, as described in (Denny et al., 2018).

Given the overall weaker binding of mutant PUM1 (Figure S6C,D), and to allow more comprehensive comparisons of single mutant binding by wild-type and mutant protein (Figure 5D), we relaxed the affinity filter for mutant PUM1 data. Rather than applying a ΔG < −8.5 threshold, we included variants with at least 15% of RNA bound by PUM1 mutant at the highest probed protein concentration.

### Assessing reproducibility and combining experimental replicates

Two replicate experiments of PUM2 binding were combined by calculating the error-weighted mean:

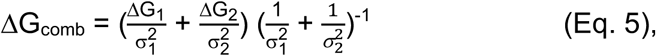

where the ΔG_1_ and ΔG_2_ are ΔG values from each replicate and Δ_1_ and Δ_2_ are 95% confidence intervals of the respective ΔG values. Weighted propagated error was calculated as:

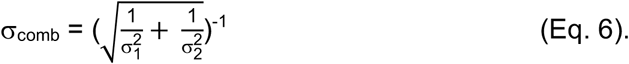

The observed small systematic offset (Figure 1E) was subtracted from Replicate 2 values before averaging to prevent distortions in cases where a variant was only present in one replicate. The offset (0.28 kcal/mol) was derived from mean replicate difference between highest affinity variants (ΔG < –9.8 kcal/mol).

### Assessing the significance of scaffold differences

The significance of single mutant scaffold differences before and after accounting for structure was assessed via a false discovery rate (FDR) approach. For each single mutant and consensus sequence, the deviation of the ΔG value (weighted replicate average, Eq. 5) from the scaffold average (ΔG_avg_) was determined and converted into a z-score:

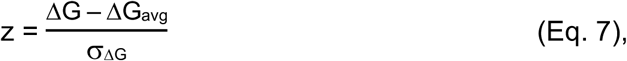

where σ is the weighted replicate error (Eq. 6). The distribution of resulting z-scores was then compared to a null distribution, where the z-scores are normally distributed around a zero mean with a standard deviation of 1. For each z-score, the number of false discoveries was determined from the probability of obtaining a value more extreme than this z-score given the null distribution (two tailed; value = 2*CDF(–|z|)), multiplied by the total number of variants. The total number of discoveries corresponded to the sum of the number of false discoveries (above) and the number of actual z-scores whose absolute values were greater than or equal to that z-score threshold. Scaffold differences were considered significant if the respective z-score had FDR ≤ 10%. Asterisks in Figures 2A, D and in Figure S2C indicate mutants where at least one scaffold showed a significant deviation from the scaffold mean.

To assess the contributions of RNA secondary structure to the scaffold variance, the above analysis was repeated using structure-corrected affinities (see **Accounting for RNA structure** below; ‘Structure-corrected’ in Figures 2B, S2A), as well as affinities of variants that lacked significant predicted structure (ΔG_fold_ > –0.5 kcal/mol; 44 of 65 total variants, including the UGUAUAUA consensus).

### Accounting for RNA structure

We used Vienna RNAfold (v. 2.1.9) (Lorenz et al., 2011) to predict ensemble stabilities for RNA structures in which the protein binding site is accessible vs. occluded due to base pairing (Figure 2C). The effect of accessibility on protein binding was defined as follows (see model in Figure S3B):

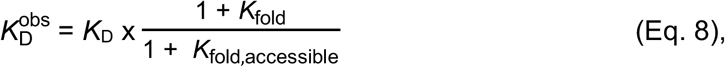

where 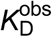 is the measured dissociation constant, *K*_D_ is the intrinsic dissociation constant (for accessible RNA), *K*_fold_ is the folding equilibrium constant that represents the ensemble of RNAs that are structured (accessible + occluded) and *K*_fold,accessible_ is the folding equilibrium constant representing RNA structures in which the protein binding site is accessible. Accessible binding sites were defined as lacking base-pairing in the 8mer core binding site (for the purpose of structure predictions) (Figure 2C). Folding equilibrium constants were based on ensemble stabilities predicted by RNAfold (`RNAfold −p0 −T 25 C` or `RNAfold –p0 –T 37 C`); stabilities of accessible RNA structures were predicted by including the constraint flag (`RNAfold –p0 –T25 C –C`) and constraining the 8mer binding site to a single-stranded state, e.g.:

UCUCUUUGUAUAUAUCUCUU

`……xxxxxxxx……,`

Where ‘x’ indicates unpaired residues. Structure effects were considered if they exceeded 0.5 kcal/mol, except as noted.

### Assessing alternative binding registers in single mutant variants

To determine potential alternative binding registers in our single mutant constructs, we computationally scanned the full RNA sequence (including scaffold and variable region) with the additive consecutive model, wherein the predicted relative affinity (expressed as ΔΔG) at each 8mer site corresponds to the sum of effects of each individual residue:

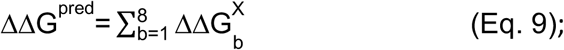

b is the position in the 8mer RNA binding sequence and X is the identity of the residue at position b. The 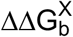 values were derived from weighted scaffold averages for each single mutant relative to the weighted average of consensus affinities, after accounting for structure effects as described in **Accounting for RNA structure**. In calculating the predicted affinities for each register, structure effects were estimated by individually constraining the respective 8mer site to the single-stranded state.

Figure S2F,G and Table S1 show the results of the initial assessment of register shifting, based on scaffold averages of mutant effects in the 5U background. No additional variants with shifted registers within less than 1 kcal/mol from the designed register were identified when the analysis was repeated for single mutants across 5A/C/U backgrounds, using mutant effects averaged across scaffolds and 5A/C/U backgrounds in Eq. 9 (Figure S3A,B).

### Development, testing and evaluation of thermodynamic binding models

To ensure the highest accuracy of model testing and global fitting, we filtered the library to only include variants without significant predicted secondary structure (ΔG_fold_ > –0.5 kcal/mol). For variants in the S1a and S1b scaffolds, we considered the ensemble structure of the entire RNA construct; for S2a and S2b scaffolds, only the stability of structures within the hairpin loop was considered, due to the high stability of the stem. The ensemble stability of structures involving the loop region was determined as the difference between RNAfold-predicted stabilities with and without the loop region constrained to the single-stranded state (ΔG_fold,loop_ = ΔG_fold_ – ΔG_fold,ss_). Only the loop sequences were used for binding predictions in global fitting.

To reduce potential systematic bias in the ΔΔG values (ΔΔG = ΔG – ΔG_WT_) that may affect the fit, we did the following. The ΔΔG values (observed and predicted) were defined relative to the UGUAUAUA<u>U</u> reference sequence rather than the slightly more stable UGUAUAUA<u>G</u> sequence (Figures 2E, S2D), because UGUAUAUA<u>U</u> was more highly represented in our library and was predicted to have less residual structure (n = 183 for ΔG_fold_ > –0.2 kcal/mol). To determine the median consensus affinity, we applied the more stringent structure cutoff of –0.2 kcal/mol (vs. −0.5 kcal/mol) due to the greater, asymmetric spread of values observed when the –0.5 kcal/mol threshold was used, consistent with residual structure effects.

Importantly, in testing and globally fitting the models below, we accounted for all possible binding modes and registers, because the strongest binding can arise from a site downstream of or partially overlapping with the original designed site, and these altered registers become more probable the larger the destabilization from mutations within the original consensus site (see Figure 4). The presence of multiple binding modes of similar affinity will also increase the overall observed affinity.

#### Testing the additive consecutive model

Additive consecutive model predictions were calculated using the equation in Figure 3A for every 9mer register in each oligonucleotide in the library, based on measured single mutant penalties (Table S2). The ensemble affinity for a given oligonucleotide was determined as illustrated in Figure 4 (‘Consecutive’; *top left*):

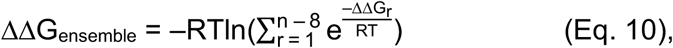

where n is the length of the oligonucleotide variant, r is the index of an individual 9mer register, ΔΔG_r_ is the predicted penalty for binding in register r, R is the gas constant and T is temperature (25 °C).

Global fitting to the additive consecutive model (Figure S4B,C) was performed as described in the *Global fitting* section below, using terms for bound residues only.

#### Base flipping analysis

For the initial assessment of the additive nonconsecutive model that permits base-flipping, we focused on oligonucleotides in our library that contained C-insertions at various positions of the consensus sequence (UGUAUAUAU). For comparisons to the additive consecutive model predictions in Figure 3C–E, we used variants without predicted register shifts to favorable sites involving flanking sequences, as this allowed for the most accurate estimation of the lower limit of the flipping penalty at the indicated site. The construct sequences used are indicated in Table S4.

#### Global fitting

To fit the final model (additive nonconsecutive with coupling; Figure 3G), the ΔΔG values for all registers with no flips, all registers with single flips, all registers with double flips, and all registers with two single nucleotide flips were included when computing the partition function for binding (see Figure 4). Registers with more than two flipped residues were not included in the partition function calculation because their predicted affinity was low enough that it minimally affected the overall binding affinity of the ensemble of bound states. The ΔΔG values for individual registers were used to compute the partition function and the overall ΔΔG for the ensemble of bound states for each RNA variant, where i is the register number (Figure 4)). This ensemble ΔΔG was compared to the experimentally observed ΔΔG values during fitting. The model assumed a single protein bound to each RNA variant, which was supported by the tight distribution of *f*_max_ values and our ability to detect binding of multiple bound protein monomers based on proportional increase in fluorescence (Figure S1E; W.R.B. et al., in preparation). The additive consecutive and additive nonconsecutive models (Figure 3A,F) were fit analogously, but with the flipping and coupling terms (additive consecutive model) or coupling terms (additive nonconsecutive) excluded from the model.

During fitting, the only values that were not allowed to vary were the bound parameters for the consensus ‘UGUAUAUAU’ residues, which were set to a penalty of 0 kcal/mol. The other bound parameters for non-consensus residues were allowed to vary within the higher of the 95% confidence interval determined from the individual single mutant measurements or ±0.4 kcal/mol from the median ΔΔG values from the individual single mutant measurements (Figure 2E, Table S2). Single and double flip parameters were essentially unconstrained during fitting and were allowed to vary between 0 and 7 kcal/mol. The two coupling terms were constrained to between –4 and 0 kcal/mol. The coupling terms were included in both consecutive and flipped registers, with the condition that no flips interrupted the series of flipped residues. The script used for global fitting can be found on https://github.com/pufmodel.

All models were initially trained on a randomly selected subset of half the data and tested on the remainder of the data to prevent overfitting. In all cases, the model performed nearly identically on the training and test sets. To compute the final parameters, the model was fit with all sequences meeting the structure cutoff. All models described in the text were fit by minimizing the sum of the squared error between the predicted and measured ΔΔG values for each sequence. To help ensure that the fit was finding a global minimum, both the BFGS and differential evolution algorithms implemented in the lmfit module in Python were used for fitting. Additionally, fits with flipping parameters were initialized to different values between 1 and 4 kcal/mol and led to the same fit parameters, providing additional support for convergence to a global minimum.

We assessed the stability of the final fit parameters (additive nonconsecutive + coupling model; Table 1, Figure 3C) by performing multiple fits with bootstrapping and a parameter sensitivity analysis. Bootstrapping was performed as follows: the 5206 sequences were sampled with replacement and the model parameters were fit to each resampled dataset. The 95% confidence intervals from the bootstrapping analysis are reported in Table S9. To examine how well bounded the model parameters were, we computed the sensitivity of the RMSE of predicted vs. observed affinities to variations in each individual parameter while holding all others constant at their fit values. Each value varied within the constraints that were applied during fitting. For some flipping parameters, the same RMSE value resulted from all parameter values greater than a given value, implying that for these parameters the penalty must be larger than a certain value so that it will not occur in the most stable register in any of our constructs, but because the parameter is so destabilizing we can only say that it must be at least as perturbative as the minimum value resulting in a constant RMSE. As a result, we reported the lower bound for these parameters (Table 1B).

#### Coupling analysis

To assess positional coupling, we tested double mutants of the UGUA[A/C/U]AUAU reference sequence for systematic deviations from predictions by the additive consecutive model. To obtain the library variants for this analysis, we filtered our mutant library for all sequences that featured a single dominant consecutive register (i.e., with less than twofold, or 0.4 kcal/mol, further stabilization provided by other consecutive or flipped registers, as predicted by the additive nonconsecutive model in Figure 3F). Variants deviating from the consensus sequence at two positions were identified and their predicted affinities were calculated by adding the respective experimentally determined single mutation penalties (ΔΔG_pred_ = ΔΔG_1_ +ΔΔG_2_; Figure 2E, Table S2). We used the experimentally derived instead of globally fit single mutant penalties in this analysis, as coupling may affect the fit values. Qualitatively, the conclusions were not affected by the fit parameters, as in both cases the strongest coupling was observed between positions 7 and 8, with negligible deviations at other positions. Only double mutant combinations represented by more than one sequence in our library were considered. Deviations between the observed and predicted ΔΔG value were determined and averaged across all mutants with mutations at a given pair of positions (Figure S4G).

The double mutant analysis indicated coupling between mutated positions 7 and 8, with negligible deviations from additivity at other positions for which data were available. Because of the highly destabilizing effects of mutations in the 5′ half of the binding site, these mutations were strongly underrepresented among double mutants, as they generally lead to alternative registers being preferred or fall outside the reliably measurable affinity range. For the same reasons, any couplings involving 5′ mutations are unlikely to be biologically relevant.

To determine the sequence dependence of position 7 and 8 couplings, including potential longer-range couplings, we next extended the analysis to varying combinations of residues flanking each position 7 and 8 residue. Specifically, and recognizing that coupling is most likely to occur between neighboring residues, we assessed the following combinations for systematic deviations from predicted values: 1) residue of interest flanked by two consensus residues; 2) only the 5′ neighboring base mutated; 3) only the 3′ neighboring base mutated; 4) both neighboring residues mutated. In this analysis, we included all variants that contained the indicated combination in the best predicted consecutive register (as predicted by additive nonconsecutive model; Figure 3B); any sequence was permitted outside the indicated combination.

Only 7G and 7C showed strong systematic deviations from predicted affinity for the indicated neighbor combinations (Figure S4H,I), and to a lesser extent––the 9G residue, which showed systematically tighter binding when preceded by the consensus residue 8A as opposed to residue 8 mutations (Figure S4J). Further inspection of 7G coupling indicated an additional bifurcation based on position 5 identity, indicating longer-range coupling (Figure S4H). The previously observed structural differences in purine and pyrimidine recognition at position 5, with dramatic effects on backbone configuration and in some structures––on position 6 recognition provide a potential structural rationale for this longer-range coupling (Lu and Hall, 2011).

### Analysis of in vivo crosslinking data

#### Determining a set of position weight matrices to find putative binding sites

The analysis of in vivo crosslinking data was carried out in two stages: first, we identified the transcriptome sites predicted to be bound by PUM2 (ΔΔG_pred_ ≤ 4 kcal/mol) by using a set of position weight matrices (PWMs) that approximate our thermodynamic model to efficiently identify binding sites. Second, we applied the full thermodynamic model, as described in Figure 3G and Figure 4, to this set of putative binding sites. Using PWMs for genome-wide searches is supported with currently available software and can computationally obtain matches to the whole transcriptome within a reasonable amount of time. In contrast, genome-wide application of the full thermodynamic model was prohibitively computationally expensive.

Each PWM is a matrix with rows representing different positions within the binding sequence and columns representing the four bases that could be at that position. The value of the PWM for each position and base is the probability of observing that base at that position in a set of binding sequences, wherein the probability is proportional to the ΔΔG terms from our thermodynamic model, as detailed below: *p*_*i,j*_= Σ_j_*exp*(ΔΔ*G*_*i,j*_/*RT*)/exp(ΔΔ*G*_*i,j*_/*RT*) for position i and base j. For the simplest binding configuration (i.e., 9 bound positions with no flipped residues), ΔΔG_i,j_ values corresponded to the binding terms in Table 1A (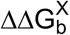). Each genomic sequence was compared with the PWM to determine a log-odds score: *s* = ∑*_i_ I_i,j_log*(*p*_*i,i*_/0.25), where *I*_*i,j*_ = 1 if the sequence at position i has base equal to j, otherwise *I*_*j,j*_ = 0. A log-odds score of ≥2 was found to capture the large majority of variants with ΔΔG < 4 kcal/mol, and so this value was applied as the threshold above which a sequence was considered as a putative binding site.

To account for binding registers with a single flipped residue, a row was inserted at the flipped position, with values derived from the base flipping penalties 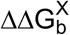, Table 1B). Our thermodynamic model found that flipping is accommodated only in four positions; thus, a PWM was determined for each of these four binding registers. In practice, for a given sequence, flips between positions 4/5 and 5/6 had very similar PWMs, and so a single PWM was derived to search for both of these binding configurations, using the average values. On average, an inserted residue penalized the overall affinity by ∼1.5 kcal/mol. To account for this overall destabilization, the threshold log-odds score for these three flipped PWMs was increased to 4 (i.e., a sequence had to have log odds score ≥4 for the sequence to be considered as a putative binding site).

A set of four additional PWMs were derived to account for having two flipped residues at each of the four flipped positions. Once again positions 4/5 and 5/6 produced very similar PWMs and were averaged, resulting in three distinct PWMs. For these PWMs, the threshold for log-odds score was increased to 5 to account for the additional destabilization of having two flipped residues.

The PWM for no flipped residues and with one flipped residue had two or one fewer positions to account for, respectively, than the PWM for two flipped residues. Thus, all PWMs were brought to the same length of 11 rows (corresponding to 11 bound or flipped positions) by padding at the 3′ end with rows that do not contribute to the log odds score of any sequence (values were set equal to 0.25 for all bases).

Finally, these seven PWMs and their respective threshold values were saved in a single file, with format defined by the program HOMER, as described: http://homer.ucsd.edu/homer/motif/creatingCustomMotifs.html.

#### Determining binding sites within the transcriptome

The seven PWMs described above were used to determine putative binding sites across the transcriptome for subsequent quantitative analysis with our full thermodynamic model. Initially, a set of genome locations was determined from the Gencode v24 annotation file, obtained from the ENCODE project:

`wget` http://www.encodeproject.org/files/gencode.v24.primary_assembly.annotation/@@download/gencode.v24.primary_assembly.annotation.gtf.gz.

These genome annotations were converted to a bed file format, and any overlapping regions were merged, using the program bedtools merge (v2.25.0). The package HOMER (v4.8.3) was used to search for matches to the PWMs within these genome locations, with the command:

`annotatePeaks.pl ${genome_locations} hg38 –m ${pwm_file} –mbed {output_motif_locations} –noann –nogene.`

The set of output motif locations, corresponding to putative binding sites, was subsequently filtered to remove any overlapping regions (bedtools merge), and if two regions overlapped, only the site with the lowest log odds score was kept. This set of motif sites was annotated again using HOMER to map each site to a gene, mRNA location (i.e. 5′ UTR, CDS, 3′ UTR), and gene type (i.e., protein-coding, noncoding RNA, etc), each of which come from the default Refseq annotations for human genome assembly GRCh38 (hg38) (O’Leary et al., 2016).

`annotatePeaks.pl ${motif_locations} hg38 > ${output_annotations}`

These annotations were used to filter motif locations based on: (1) Being part of a protein-coding gene in the 5′ UTR, CDS, 3′ UTR, or being part of a noncoding RNA; (2) Being on the same strand as its annotated gene. This filtering resulted in a final set of 646,268 binding motif locations around which PUM2 binding was assessed.

In addition to this set of filtered motif locations, a set of “random” sites were determined, which served as controls throughout. These sites were obtained by choosing 5,000,000 random start sites within the original set of genome locations. These random sites spanned the same number of nucleotides as the putative binding sites (11 nt). These 5 million sites were subsequently annotated and filtered exactly as the putative binding sites were, resulting in 76,094 “random” locations.

### Using the thermodynamic model to predict ΔΔG values of each putative binding site

The 11-nt sequence of all motif locations (“binding” or “random”) were each assessed for their predicted ΔΔG using the full thermodynamic model (Figure 4; 37 °C). The sequence within each motif location was determined using the bedtools getfasta command and the hg38 genome build in fasta format.

This 11-nt window has three possible binding registers if no residues are flipped (ΔΔG_consecutive,1_,ΔΔG_consecutive,2_, ΔΔG_consecutive,3_), in addition to other binding registers with one or two residues flipped (see Figure 4). Each sequence was also assessed for the ΔΔG_noflip_, which represents the ensemble energy of the three ΔΔG_consecutive_ values, with no contribution from any of the flipped binding registers.

### Determining expression of putative PUM2 binding sites

RNA-seq expression data for K562 cell lines was obtained from the ENCODE project (Consortium, 2012). These data consisted of transcript-per-million (TPM) values for each ENSEMBL transcript across two replicates (https://www.encodeproject.org/files/ENCFF272HJP/@@download/ENCFF272HJP.tsv and https://www.encodeproject.org/files/ENCFF471SEN/@@download/ENCFF471SEN.tsv). The TPM values for each transcript were averaged across the two replicates, and the value for each Refseq transcript identifier was then determined using Ensembl Biomart for hg38 (http://uswest.ensembl.org/biomart/martview/2602e63453effc6fdd3fae6b78dfed42). The TPM value for each Refseq transcript gave the relative expression of motif sites within that transcript, as assessed using the annotations from HOMER described above.

In some analyses, only motif sites on transcribed genes (TPM > 0 in K562) were included in certain subsequent analyses, resulting in further filtering of the number of motif sites examined to 445,368 total sites.

### Obtaining eCLIP signal around putative PUM2 binding sites

Enhanced UV crosslinking and immunoprecipitation (eCLIP) data for PUM2 protein in human K562 cells was obtained from ENCODE (Van Nostrand et al., 2016). Sequencing read alignments (in the form of a BAM file) were obtained for two replicate pulldown samples and one input sample which did not undergo antibody pulldown for PUM2 but was otherwise experimentally processed identically (https://www.encodeproject.org/files/ENCFF786ZZB/@@download/ENCFF786ZZB.bam, https://www.encodeproject.org/files/ENCFF732EQX/@@download/ENCFF732EQX.bam, https://www.encodeproject.org/files/ENCFF231WHF/@@download/ENCFF231WHF.bam). Only alignments corresponding to the second sequencing read (read2) were kept, as the 5′ end of this read corresponds to the putative crosslinking site (Van Nostrand et al., 2016). The alignments were obtained using the package samtools (version 0.1.19-96b5f2294a):

`samtools view –bh –f 128 ${input_bam} > ${output_R2_bam}`

This set of filtered alignments was used to determine the number of observed crosslinks (i.e., read2 start sites) on each strand at any position within the genome, using the package bedtools:

`bedtools genomecov –ibam {R2_bam} –strand + –bg –5 > {output_bedgraph_plus}`

`bedtools genomecov –ibam {R2_bam} –strand – –bg –5 > {output_bedgraph_minus}`

The number of crosslink sites (from eCLIP data) at each nucleotide position within each motif location was determined using the package pyatac ins (https://nucleoatac.readthedocs.io/en/latest/pyatac/). Only crosslink sites on the same strand as the motif were included in the total count. The number of reads starting 55 nt upstream and extending to 25 nt downstream from the motif location center (80 nt total; Figure S7F) were summed for each sample, corresponding to the motif site’s eCLIP read count; this window accounted for the asymmetrical observed distribution of crosslink sites around PUM2 consensus motifs. eCLIP signal for each motif site was determined as the sum of eCLIP read count for the two replicates, divided by the relative expression of that motif site. Similarly, eCLIP input for each motif site was the eCLIP read count for the input sample, divided by the relative expression of that motif site.

The median eCLIP signal and input around sites identified as ‘background’ sites was determined from sites with predicted ΔΔG > 4.5 kcal/mol, regardless of whether the site originated from the “binding” or “random” motif locations. The eCLIP signal (or input) values were divided by the ‘background’ signal (or input) value to obtain the eCLIP signal (or input) enrichment above the background expectation.

### Assessing secondary structure around motif sites

Consensus motif sites (ΔΔG_pred_ < 0.5 kcal/mol), comprising 4,816 non-overlapping sites, were assessed for local secondary structure occluding the binding site. The sequence around each motif site was determined using `bedtools getfasta`. Multiple different lengths of flanking regions were included in this assessment (10 nt, 20 nt, or 80 nt on either side of the motif site). An additional line within the sequence fasta file gives the constraint that the binding site (i.e., the first 8 nt of the motif site, given the weak interaction at position 9) remains unpaired. Using the program RNAfold (v2.1.8 (Lorenz et al., 2011)), the ensemble energy was determined for each sequence:

`cat ${input_fasta} | RNAfold –-noPS –p0 –C –T37 > ${output_values_wconstraint}`

`cat ${input_fasta} | RNAfold –-noPS –p0 –T37 > ${output_values_noconstraint}`

The difference in ensemble free energy with and without the constraint gives the accessibility of that site: ΔΔG_ss_ = ΔG_no_constraint_ – ΔG_constraint_ (see also **Accounting for RNA structure** above).

### Enrichment of PUM2 sites within 3′ UTRs

Sites derived from the PUM2 PWMs (described in **Analysis of in vivo crosslinking data** above; not filtered for being expressed in K562 cells) were divided into bins based on the predicted ΔΔG, and the fraction of motifs within each bin that were annotated as 5′ UTR, CDS, or 3′ UTR was determined as described above (i.e., using HOMER), and are shown in Figure 6C. Enrichment for transcript annotations of motif sites (Figure S7E) was determined relative to the annotation frequencies of “random” locations.

### HPLC purification of RNA oligonucleotides for competition binding measurements

Desalted RNA oligonucleotides were ordered from IDT and purified by reverse-phase HPLC (XBridge Oligonucleotide BEH C18 Prep column or Agilent ZORBAX Eclipse Plus C18 column), using an acetonitrile gradient in the presence of 0.1 M triethylamine acetate. Following purification, the solvent was exchanged into MilliQ water with Amicon Ultra 3K concentrators.

### [γ- ^32^P]-labeling of RNA nucleotides

RNA oligonucleotides for direct binding measurements were ordered from IDT and 5′ labeled with [γ- ^32^P] ATP (Perkin Elmer) using T4 polynucleotide kinase (T4 PNK, Thermo Fisher Scientific). The 5 μL reactions contained 1x PNK buffer (Thermo Fisher Scientific), 5 μM oligonucleotide, 5 μM [γ- ^32^P] ATP and 1 μL of T4 PNK. The reactions were incubated at 37 °C for 30 min and purified by non-denaturing gel electrophoresis (20% acrylamide).

### Gel shift binding measurements

#### Competition binding measurements

To obtain PUM2 binding measurements in the absence of potential structure formation and alternative sites, and to compare the affinities determined by different approaches, we performed competition gel shift binding measurements with 8mer oligonucleotides carrying a subset of single mutations in the UGUAUAUA background. PUM2 (0.68 nM) was combined with trace labeled “S1a” RNA (UCUCUU<u>UGUAUAUA</u>UCUCUU, <0.08 nM) in binding buffer (2 mM DTT, 100 mM potassium acetate, 0.2% Tween-20, 20 mM sodium HEPES, pH 7.4, 5% glycerol, 0.1 mg/mL BSA (NEB), 2 mM MgCl_2_), and diluted two-fold into solutions containing varying concentrations of unlabeled competitor RNAs (3-fold serial dilution series; 7–8 concentrations per oligonucleotide; final concentrations of 0.34 nM PUM2, <0.04 nM labeled S1a RNA, 0.17––3330 nM competitor RNA, depending on the oligonucleotide). Binding reactions were incubated at 25 °C for at least 1 h; equilibration was established by measuring binding after 1 h and 4.5 h incubations, which gave consistent results. We also performed controls for titration effects, by incubating the most tightly bound oligonucleotides (consensus, 5G and 7G variants) with 0.16 or 0.32 nM PUM2 (final concentration), giving consistent affinities. Following equilibration, 7.5 μL aliquots were transferred to 7.5 μL ice-cold loading buffer (5% Ficoll PM 400 (Sigma), 0.03% BPB, and 2 μM unlabeled S1a RNA in binding buffer). The low temperature and unlabeled consensus RNA in the loading buffer prevented changes due to potential re-equilibration during sample loading (Vaidyanathan et al., 2017)). The samples were carefully and immediately loaded on a continuously running 20% native acrylamide gel (5 °C, 750 V, 0.5x Tris/Borate/EDTA (TBE) running buffer: 44.5 mM Tris-borate, 1 mM Na_2_EDTA, pH 8.3; DANGER: extreme caution is required in this step due to high voltage; https://ehs.stanford.edu/reference/electrophoresis-safety). Gels were dried, exposed to phosphorimager screens and scanned with a Typhoon 9400 Imager.

Binding affinity for the labeled S1a oligonucleotide was measured in parallel by incubating 0.0038–81 nM PUM2 (3-fold serial dilutions) with trace labeled S1a RNA (<0.04 nM) in binding buffer for at least 1 h at 25 °C. Samples were analyzed by gel electrophoresis as above. Measurements with three labeled RNA concentrations across a 9-fold range (upper limits of 13– 120 pM) gave consisted results, indicating no titration effects. Sufficient equilibration time was established by measuring the dissociation rate (0.011 s^−1^, corresponding to 5.25 min upper limit of equilibration time––i.e., five half-lives; see below (Vaidyanathan et al., 2017)).

The gels were quantified with TotalLab Quant and fitting was performed with KaleidaGraph 4.1 (Synergy). The affinity for the labeled S1a RNA was determined by fitting to a single-site binding equation:

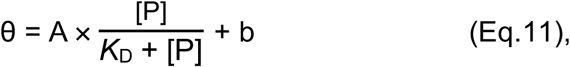

where θ is fraction bound RNA, A is amplitude, [P] is PUM2 concentration, *K*_D_ is the equilibrium dissociation constant and b is background. Competitor affinities (*K*_D,comp_) were determined using the equation by Lin & Riggs (Lin and Riggs, 1972):

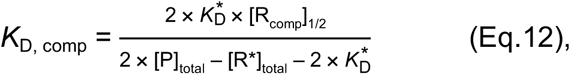

where K_D_* is the dissociation constant of the labeled S1a RNA; [R_comp_]_1/2_, the competitor concentration at which half of the labeled RNA is bound; [P]_total_, the protein concentration; and [R*]_total_ the labeled RNA concentration. To determine the fraction of competitor RNA at which half of labeled RNA was bound, the competition binding curves were normalized by the fraction of labeled S1a RNA bound at saturation with no competitor (0.94). [R*]_total_ was the upper limit of the labeled RNA concentration based on the total input and elution volume in the labeling reaction (<0.04 nM). Using the lower limit based on scintillation measurements of the ^32^P label (∼0.004 nM) did not affect relative affinity calculations and affected absolute affinities by <10%. The values shown in Figure S3F,G are averages and standard errors from two replicate measurements.

For determination of flanking sequence effects, CUUGUAUAUAN oligonucleotides (N = A/C/G/U) were ordered from IDT, 5′ end labeled with [γ- ^32^P] ATP, and binding was measured as described for the S1a RNA above.

#### Dissociation rate constant measurements

PUM2 dissociation rate constant from S1a RNA was measured by incubating 3.8 nM PUM2 with labeled S1a RNA (<0.5 nM) in binding buffer at 25 °C for 50 min, followed by addition of 2.5-fold volume excess of unlabeled RNA chase in binding buffer (final concentrations: 1 nM PUM2, <0.14 nM labeled S1a RNA, 1 μM unlabeled S1a RNA). At various time points, 7.5 μL aliquots were moved to 7.5 μL ice-cold loading buffer (5% Ficoll PM 400 (Sigma), 0.03% BPB in binding buffer) and immediately loaded on continuously running 20% native acrylamide gel. The dissociation curve was fit to a single exponential in Kaleidagraph:

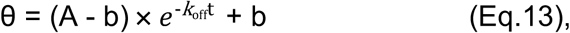

where θ is fraction bound RNA, A is the fraction bound before adding the chase (A = 0.90), b is the fraction bound at the completion of the dissociation reaction (b = 0.02), and *k*_off_ is the dissociation constant, and t is time after adding chase.

#### Determination of active protein fraction by titration

Saturating concentration of unlabeled consensus RNA (10–200 nM; S1a or UCUUGUAUAUAUA for wild-type PUM1/2, UCUUGUAUUUAUA for mutant PUM1) was mixed with trace ^32^P-labeled RNA of the same sequence (<0.15 nM) and incubated with protein concentrations at least 4-fold below and above the RNA concentration for 45 min – 1 h (25 °C). Following native gel electrophoresis, active protein fraction was determined from the intersection of lines fit through protein concentrations below and above the breakpoint. Throughout, the protein concentrations and absolute affinities reflect active protein concentrations (SNAP-Cy3B-PUM2: 57%, PUM1-SNAP-Cy3B: 61%, mutPUM1-SNAP-Cy3B: 20%, unlabeled SNAP-PUM2 used for gel-shift controls: 38–45%).

### Modeling the cellular PUM2 binding landscape

To assess the distribution of PUM2 across cellular RNA sites, we determined the numbers of each 9mer, 10mer and 11mer sequence in the human transcriptome (representing consecutive sites and sites containing one or two flipped residues). For simplicity, here we assumed equal expression of all protein-coding transcripts, with sequences obtained from GENCODE (genome release GRCh38.p12; ‘Protein-coding transcript sequences’ fasta file; **<u>Error! Hyperlink reference not valid.</u>**et al., 2012). Absolute numbers of each binding site were determined by normalizing the nucleotide count in the above transcriptome file to match the estimated mRNA nucleotide count in a single human cell (8.9**×**10^8^ nucleotides, corresponding to ∼0.5 pg mRNA per cell) (Livesey, 2003; Marinov et al., 2014; Tang et al., 2011). These numbers can be adjusted to account for cell-specific variation in total mRNA levels and differential expression based on publicly available RNAseq data (Consortium, 2012; Sloan et al., 2016).

The number of PUM2 molecules per cell was estimated at 10,000, based on published numbers of 2,000 and 18,000 in HCT116 and HeLa cells, respectively (Lee et al., 2016; Nagaraj et al., 2011).

To calculate the distribution of cellular protein bound across the different mRNA sequences, we first calculated PUM2 relative affinities for each 9–11mer site using our thermodynamic model. The affinities for consecutive 9mer sites were calculated using the binding and coupling terms in Table 1A,C; to determine affinities for sites containing flipped residues, we calculated the ensemble affinities of the four possible registers with one single-nucleotide flip (Figure 4; 10mer sites); and the ensemble of the four possible registers with a two-nucleotide flip and six possible registers of two single-nucleotide flips (11mer sites), using the terms in Table 1A–C.

PUM2 occupancies for each RNA species were calculated using an equilibrium competition model, where the occupancy of a given RNA species (P*Ri) is a function of protein abundance (P), and the affinities and abundances of all RNA (R) sites:

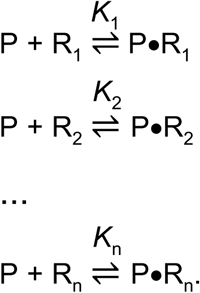

The fraction of total protein bound to RNA sequence R_1_ equals:

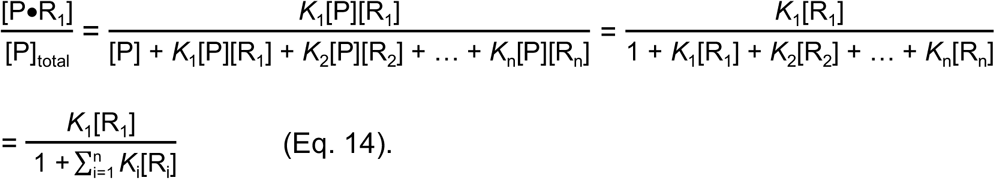

We used affinities predicted for each RNA by our thermodynamic model:

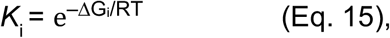

where ΔG_i_ = ΔG_WT_ + ΔΔG_i_; ΔG_WT_ is the measured affinity for the UGUAUAUAU reference sequence at 37 °C (–12.1 kcal/mol), and ΔΔG_i_ is the relative free energy for binding to sequence R_i_ (ΔΔG_i_) predicted by our thermodynamic model.

To convert the sequence counts into concentrations, we used the cell volume of 10^−12^ L (Fujioka et al., 2006). Given the much greater number of RNA sites than the number of cellular PUM2 molecules (78,000 consensus UGUA[ACU]AUAN sites alone vs. 10,000 PUM2 molecules), we assume that most RNA sites are unbound, i.e., that [R_i_] ≈ [R_i_]_total_. [This assumption will not hold at very high protein concentrations or in the presence of very high specificity, as discussed in main text; these alternate regimes can be readily simulated using KinTek Explorer or similar software (Johnson et al., 2009).]

The occupancies plotted in Figure 6G correspond to the amount of PUM2 bound to each RNA species (i.e., fractional protein occupancy from Eq. 14 multiplied by the total amount of protein):

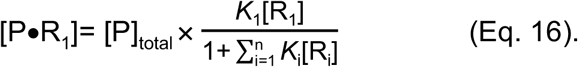

Concentrations of occupied sites were converted into the number of protein bound sequences as follows: [P•R_1_] **×** *N*_A_ × V_cell_, where *N*_A_ is Avogadro’s number and V_cell_ is the cell volume.

The bars in Figure 6G denote the sum of occupancies for all 9mer RNA species containing the indicated numbers of nonconsensus residues (blue) and 10–11mer species with flipped residues (green).

Fractional occupancies of each RNA sequence were determined (using the definition of the amount of bound RNA from Eq. 14), as follows:

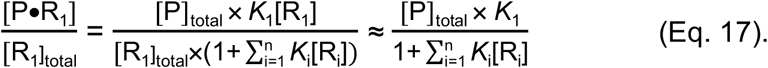

Given the varied affinities for RNAs containing a given number of nonconsensus residues, there is a range of fractional occupancies, as depicted in the boxplot in Figure 6I.

#### Occupancy prediction algorithm

A script for predicting PUM2 occupancy on any RNA sequence (see Figure 6E,F) can be found on https://github.com/pufmodel. The script accepts any RNA sequence in fasta format and predicts PUM2 occupancies relative to the UGUAUAUAU consensus at each site along the sequence based on our thermodynamic model. The algorithm currently assumes a linear relationship between affinity and occupancy, as explained above in **Modeling the cellular PUM2 binding landscape**, as we do not expect saturating binding in vivo in the presence of the large excess of tight RNA binding sites over cellular protein. Currently the script provides normalized occupancies relative to the consensus sequence; to determine fractional RNA occupancies within the cell, experimental PUM2 and RNA concentrations will need to be considered. The fractional occupancies in the example shown in Figure 6F have been calculated based on our landscape model (see **Modeling the cellular PUM2 binding landscape**).

